# Multifaceted luminance gain control beyond photoreceptors in *Drosophila*

**DOI:** 10.1101/2023.03.17.533132

**Authors:** Madhura D. Ketkar, Shuai Shao, Julijana Gjorgjieva, Marion Silies

## Abstract

Animals navigating in natural environments must handle vast changes in their sensory input. Visual systems, for example, handle changes in luminance at many timescales, from slow changes across the day to rapid changes during active behavior. To maintain luminance-invariant perception, visual systems must adapt their sensitivity to changing luminance at different timescales. We demonstrate that luminance gain control in photoreceptors alone is insufficient to explain luminance invariance at both fast and slow timescales and reveal the algorithms that adjust gain past photoreceptors in the fly eye. We combined imaging and behavioral experiments with computational modeling to show that, downstream of photoreceptors, circuitry taking input from the single luminance-sensitive neuron type L3 implements gain control at fast and slow timescales. This computation is bidirectional in that it prevents underestimation of contrasts in low luminance and overestimation in high luminance. An algorithmic model disentangles these multifaceted contributions and shows that the bidirectional gain control occurs at both timescales. The model implements a nonlinear interaction of luminance and contrast to achieve gain correction at fast timescales and a dark-sensitive channel to improve the detection of dim stimuli at slow timescales. Together, our work demonstrates how a single neuronal channel performs diverse computations to implement gain control at multiple timescales that are together important for navigation in natural environments.

**Significance Statement:** Luminance varies slowly throughout the day, or rapidly during movements. Yet, animals show luminance-invariant behaviors to visual cues across these scenarios. Combining in vivo imaging, behavioral analyses and computational modeling, we here show that diverse luminance gain control operations required for robust visual processing exist in the fly visual circuitry beyond photoreceptors. Despite relying on a single luminance-sensitive neuron type, this luminance gain control is multifaceted, in that it achieves both gain increments and decrements at two different timescales. Overall, the study sheds light on a circuit gain control strategy operating at multiple timescales, which together challenge luminance-invariant visual behaviors in natural environments.

## Introduction

In natural environments, visually guided animals are challenged by constantly changing viewing conditions. In any condition, vision depends on the light reflected by an object that comprises a contrast cue relative to its surrounding. However, the difference in reflected light between the object and its surrounding changes with changing illumination. This is true when the day approaches sunset, but also when the object suddenly moves into the shade. Yet, for the object to drive a stable behavioral response, perception of contrast must be invariant to different amounts of light reaching our eyes at all timescales. For example, the black and white patches of a soccer ball are always perceived as black and white, at different times of the day, or when the ball has been kicked into the shadow of a tree (Figure 1), suggesting luminance-invariant visual processing. Luminance invariance has indeed been reported experimentally in humans, and is also seen in behavioral response of fruit flies, as well as in the neuronal response in several species (1–4). To achieve invariance, visual systems adjust their sensitivity by controlling their luminance gain, thus determining the response size relative to the mean luminance. Gain control serves two major functions: First, it enables the limited operating ranges of visual neurons to deal with the vast range of environmental light intensities, avoiding saturation (5–7). Second, gain control brings the benefit of luminance-invariant encoding of contrast, for example Weber contrast. To achieve stable perception, visual systems must be able to increase or decrease gain when a decrease or increase in luminance is encountered, respectively. For example, when a soccer ball flies into a shadow, the difference in light reflected by the black and white patches goes down, and vision thus requires a higher gain to perceive a similar contrast as compared to when the ball is out of the shade. Conversely, a reduction in gain must adjust contrast signals when the ball is back in the sunlight (Figure 1). These two types of gain control must also be in place at the different timescales: when the luminance changes rapidly within a scene (contextual luminance changes, such as open field-to-shadow), or when the overall luminance of a scene is changing slowly (absolute luminance changes, such as day-to-night)(7). Which gain control mechanisms act at different timescales in these different scenarios to contribute to luminance-invariant visual behavior, remains unclear.

**Figure 1:**
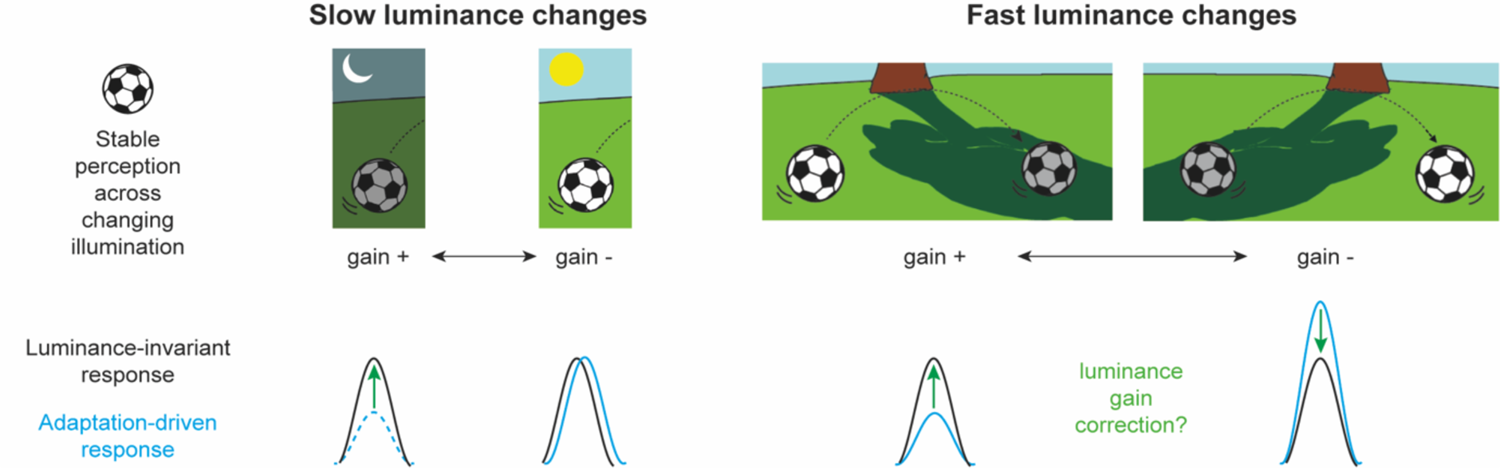
Luminance-invariant perception may require luminance gain correction past photoreceptors. Luminance gain control is necessary for stable perception of an object (here, a soccer ball) across both slow (e.g., day-night) and fast (e.g., open field-shadow) luminance changes. Gain control through adaptation may not achieve a luminance-invariant response – a response that stays the same across all luminance conditions (black). The adaptation-driven response (blue) may underestimate stimuli in dim conditions (dashed blue), in addition to under- and overestimating them across rapidly luminance-changing scenarios. Thus, gain correction (green) may be necessary at both slow and fast luminance changes.

Gain control is a characteristic of many visual neurons, from photoreceptors to cortical cells (3, 8–12). Luminance gain control implemented by photoreceptor adaptation is particularly well studied. Adapting photoreceptor types are sensitive to light changes over several orders of light intensity (3). However, photoreceptor adaptation alone might not achieve luminance invariance (Figure 1). First, photoreceptors do not adapt in low intensity ranges (3, 13, 14). Second, although the initiation of photoreceptor adaptation is fast and modulates photoreceptor sensitivity on the order of 200-300 ms (15–17), it takes tens of seconds to minutes to complete. Perfect gain control that results in the encoding of Weber contrast is achieved only at these slow timescales (3, 13, 18). Since light intensities do not only vary with slow day-night changes but also with fast movements, such as saccadic eye movements or self-motion, photoreceptor gain control alone will not establish invariance. Thus, a rapid luminance gain control mechanism must be present in visual circuitry past photoreceptors.

One mechanism for post-photoreceptor luminance gain control implementing invariance at fast timescales has recently been revealed in the fly visual system (4). Postsynaptic to photoreceptors, the lamina monopolar cells (LMCs) filter visual input differently: transient L2 response encodes changes in luminance, sustained L3 response is predominantly sensitive to luminance itself, and L1 response lies in-between with both transient and sustained response components (4, 19, 20). LMCs themselves do not achieve luminance invariance at fast timescales, but their outputs must be corrected in the downstream circuitry to ultimately achieve luminance-invariant behavior (4, 13). This corrective mechanism relies on a luminance-sensitive signal preserved past photoreceptors in L3 neurons (4, 19). A system lacking this L3-dependent luminance gain was less sensitive to dimmer stimuli when luminance was quickly changing (contextually dim stimuli), indicating that the prevailing gain was lower than required (4). While this work offered first insights into how the fly visual system handles fast luminance changes, we do not fully understand the mechanisms by which the visual system handles many different changes in the visual environment, at both slow and fast timescales.

Here, we use imaging and behavioral studies combined with modelling to identify the circuit strategies and computational principles for the fast and slow gain control occurring downstream of photoreceptors. Comparing behavior with contrast-sensitive LMC responses shows that the gain control by fly photoreceptors and LMCs is insufficient to explain behavioral invariance at both fast and slow timescales. By genetically blocking a single luminance-sensitive LMC subtype L3, we demonstrate its key contribution to luminance gain control at both fast and slow timescales, avoiding underestimation or overestimation of contrast. Thus, the L3 pathway corrects for multiple deficits in peripheral gain control. To understand how the luminance and contrast channels interact in downstream circuitry to enable these seemingly diverse actions of L3, we implemented an algorithmic model. The model predicts that contrast and luminance signals interact nonlinearly to achieve the correction at fast timescales, whereas at slow timescales, independent excitatory and inhibitory pathways downstream of L3 confer gain increment and decrement, respectively. In sum, our study describes a versatile gain control mechanism in fly vision, implemented by a single luminance-sensitive input that serves behavioral invariance and also improves vision in extremely dim conditions.

## Results

### Fly behavior uses a slow luminance gain past the second visual synapse

Visual behaviors of many species, including flies, are luminance invariant. We first tested to what extent this is already achieved by adaptation in photoreceptors at slow changes in mean luminance, over tens of seconds or minutes. To do so, we compared fly behavioral response with neuronal response to a range of contrasts downstream of photoreceptors (Figure 2; Figure S1). We measured physiological response in the first order interneuron L2, an LMC crucial for motion-guided behavior (4, 21, 22). Behavior was measured as turning response of flies positioned on an air-cushioned ball to moving OFF edges of different Weber contrasts (−50% to −87%). The background for each edge was shown for 500 ms before the edge moved across the screen for 750 ms at 192°/s, and two edge presentations were separated by 1 s of darkness. Responses to this set of contrasts were also measured in five different luminance regimes, using neutral density (ND) filters, together encompassing five orders of luminance (Figure 2A left). Thus, different contrasts were changing fast, but because different contrast sets were measured minutes apart, we could compare slow changes between the same contrast at different luminances.

**Figure 2:**
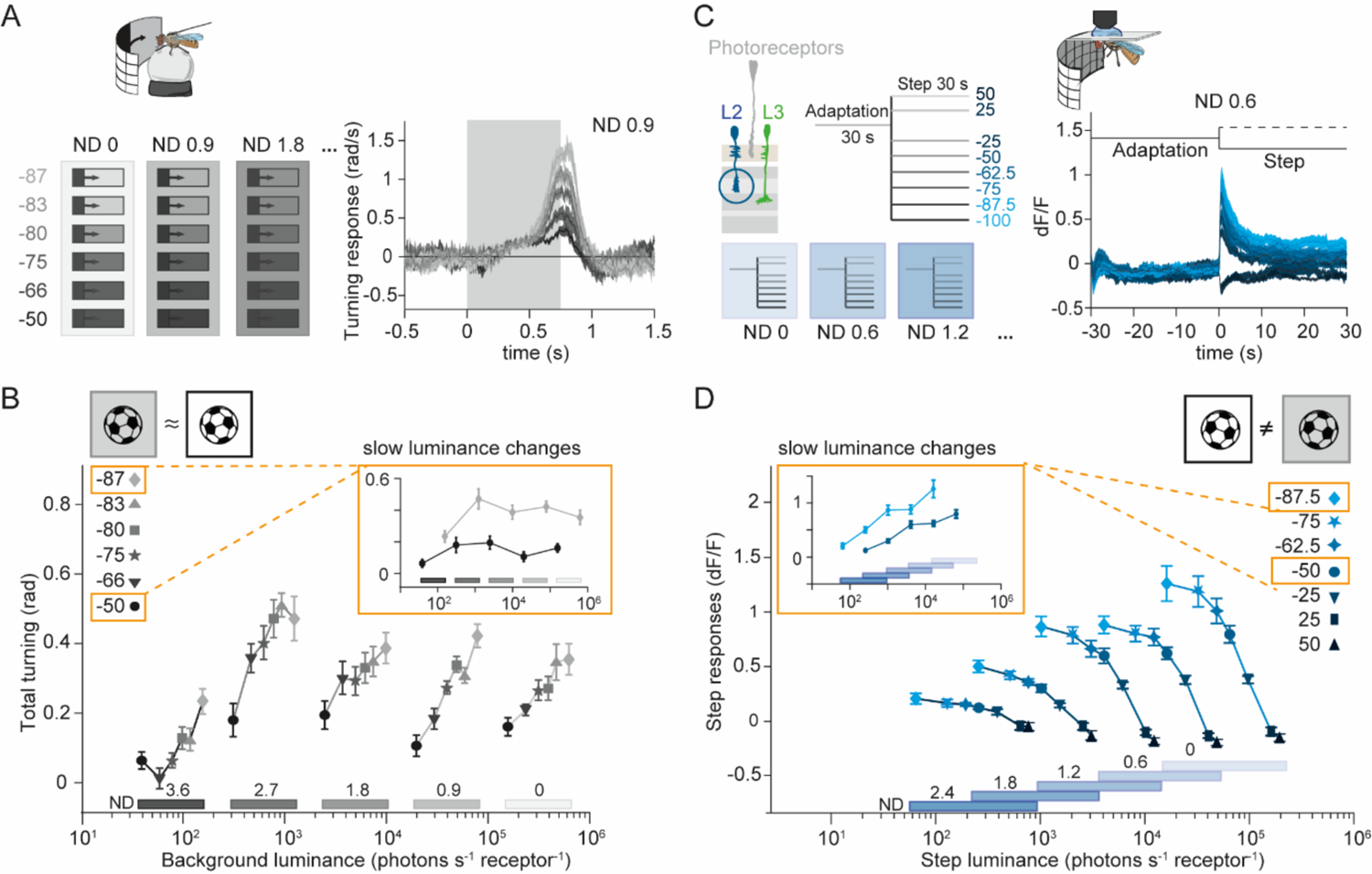
Fly behavior uses a slow luminance gain past the second visual synapse. (A) (Left) Schematic of the stimulus used to measure turning response of flies walking on a ball. Single, moving OFF edges of varying contrast were randomly interleaved within each adaptation context (denoted by neutral density (ND) 0, 0.9 etc.). (Right) Turning velocity time traces in response to different contrasts in one of the five adaptation contexts (ND 0.9). (B) Total turning in response to the contrast stimuli in all five adaptation contexts, plotted against background luminance. The inset depicts only the −50% and −87% contrast data, compared over adaptation context (i.e. slow luminance changes). Response to a specific contrast did not vary monotonically with luminance. Sample size n=10 flies per data point. (C) Schematic of the stimulus used for *in vivo* L2 calcium imaging. Within each adaptation context (denoted by ND 2.4, ND 1.8 etc.), flies were exposed to 30 s-long bright screen, followed by a 30 s-long step comprising one of eight different contrasts (% values are shown). L2 calcium response time traces in one of the luminance regimes (ND 0.6) are on the right. (D) L2 step response quantified as the difference between peak response and preceding plateau response, plotted against step luminance. The inset depicts only the −50% and −87.5% contrast data, compared over adaptation contexts (i.e. slow luminance changes). For each contrast, the response varied with luminance. Sample size is n=10 (136), #flies (#ROIs, regions of interest). Traces and plots in B-E show mean ± SEM.

Within one luminance regime, which corresponds to one adaptation context, the turning response of the fly scaled with Weber contrast (Figure 2A right). Across all but the dimmest adaptation context, turning response curves appeared to be shifted versions of each other, showing that behavioral responses to each contrast were similar (Figure 2B, Figure S1A). This was also evident when comparing the response to the same contrast taken from different adaptation contexts (Figure 2B, inset). Since the same contrast stimuli were shown minutes apart, this shows that fly behavior is luminance invariant at slow timescales. In the dimmest adaptation context, flies turned less because they likely reached sensitivity threshold. Using *in vivo* two photon imaging, calcium response of L2 neurons was also measured in five adaptation contexts (Figure 2C). Within each adaptation context, flies expressing GCaMP6f in L2 neurons were adapted to one luminance for 30 s, before luminance stepped down (6 OFF steps) or up (2 ON steps) to show one of eight values ranging from −25% to +50% Weber contrast. L2 calcium signals recorded in L2 axon terminals were negatively correlated with contrast (21) and scaled with contrast within each adaptation context (Figure 2C,D, Figure S1B). Across adaptation contexts, the L2 response curves did not appear to be shifted versions of each other, but the operating range of L2 increased with adapting luminance (Figure 2D). When examining the same contrast taken from different adaptation contexts, L2 responses did not achieve constant contrast sensitivity but retained luminance dependence, meaning that even after long adaptation times L2 responses were not fully adapted (Figure 2D, inset).

To summarize, when adapting luminance was changing slowly, we found that the turning response of flies to individual contrasts was similar across adaptation contexts (Figure 2B), whereas L2 calcium response was not (Figure 2D). Thus, behavior must utilize luminance gain control past the second order neuron L2 to achieve stable contrast sensitivity across slowly changing luminance.

### Fly behavior uses an additional luminance gain at fast timescales

The discrepancy between L2 response and fly behavior suggests that even at slow timescales (>30 s), L2 is not fully adapted. To test the timescales of L2 adaptation, we examined the temporal characteristics of L2 by fitting an exponential curve to its OFF-step calcium response (Figure 3A,B, same data as Figure 2C,D). In line with the observation that L2 response strongly varies with luminance even at slow timescales (Figure 2D), the L2 response gain rose with increasing adaptation context (Figure S2A). Moreover, most L2 responses were still decreasing at the end of the 30 s step duration, such that the steady-state L2 signals were non-zero especially in the brighter adaptation contexts (Figure 3A, Figure S2B), indicative of a slower adaptation process. Time constants of the fast adaptation process captured by exponential fits varied across luminance regimes, with the fastest time constants being in the range 1.2 s – 1.7 s, meaning that the fast adaptation processes also required more than 5 s to reach steady state (Figure 3b, right).

**Figure 3:**
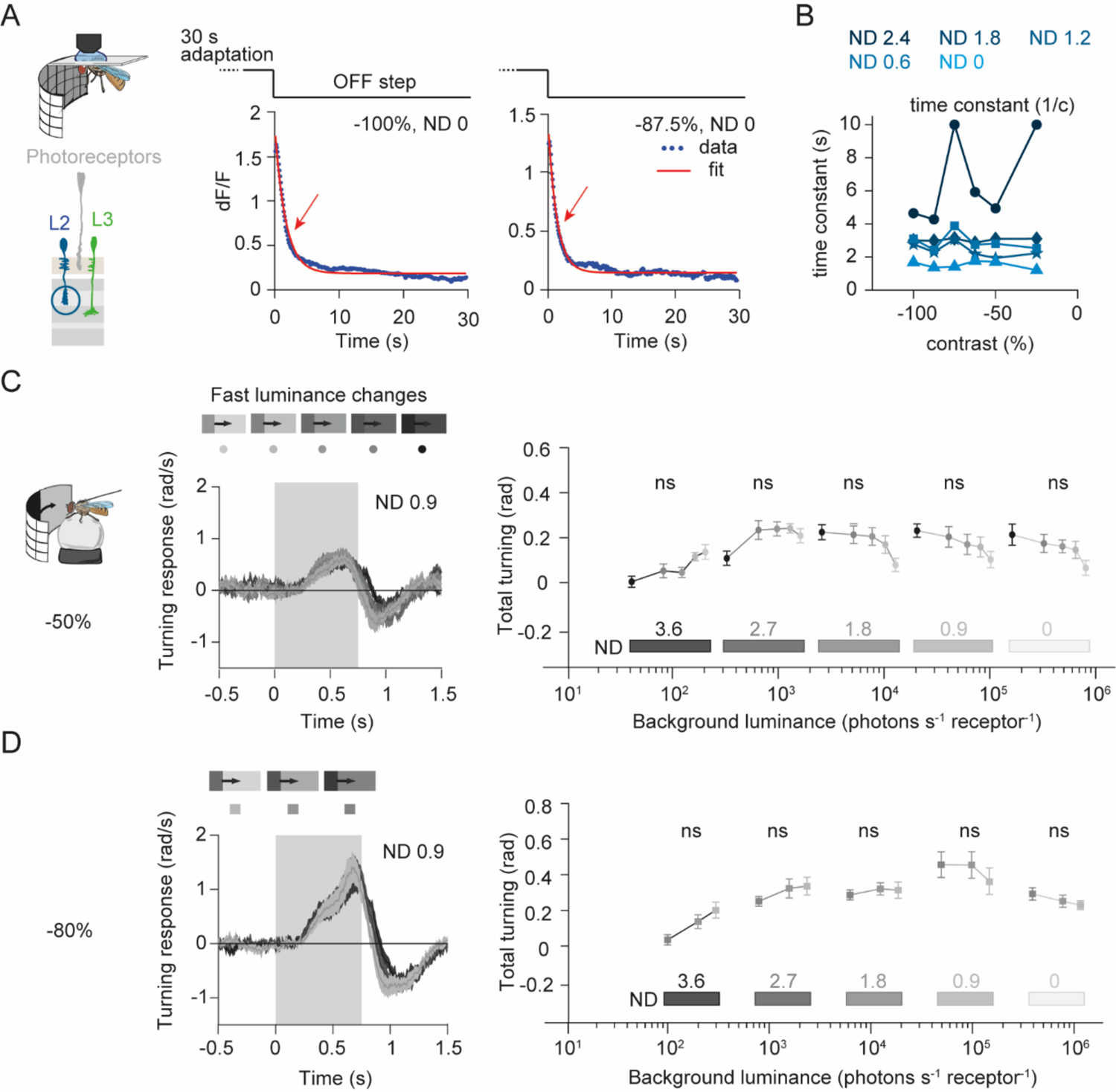
Fly behavior uses an additional luminance gain at fast timescales. (A) The mean L2 response to the −100% and −87.5% contrast OFF steps recorded at ND 0 (from the same dataset as in Figure 2D) and exponential fits to the response. Red arrows indicate the initial decrease of L2 signals arising from fast adaptation processes primarily captured by the exponential fit. (B) Time constants 1/*c* of the exponential fit (L2 response = *a* + *b* * exp(−*c* * time)) to the L2 data (all data from Figure 2D). Blue shades represent adaptation context (denoted by the ND filter). Sample size is n=10 (136), #flies (ROIs). (C,D) Turning response of flies to moving OFF edges of −50% (C) and −80% (D) Weber contrasts presented at different background luminances. (Left) Response time traces of one adaptation context ND 0.9 are shown. Within an adaptation context, 5 different luminances were used to generate −50% contrast stimuli (C) and 3 different luminances were used to generate −80% contrast stimuli (D). (Right) Total turning at all luminances and adaptation contexts. Gray shades of the symbols represent the stimulus intensity within each context. Luminance invariance was tested by one-way ANOVA corrected for multiple comparisons, ns = non-significant (indicates luminance-invariant response within one adaptation context). Sample size is n=10 flies per data point. All traces and plots show mean ± SEM.

Before reaching steady state, L2 would be even further away from achieving a constant contrast sensitivity. This suggested an additional, fast luminance gain control that scales response postsynaptically to L2. We thus tested if additional luminance gain control past LMCs contributes to the behavior at fast timescales when luminance changes within a second. For this purpose, we performed a similar behavioral experiment as above, but now kept contrast the same while changing background luminance at each stimulus presentation, allowing us to test for fast luminance gain control. We chose two contrasts – −50% and −80% – and presented them at quickly changing luminances. We observed that fly turning to one contrast was indistinguishable regardless of background luminance (Figure 3C,D). To see if this holds across a wide range of luminances, we then repeated the experiment at many different adaptation contexts. The turning response was highly similar within each of the adaptation contexts, i.e. luminance invariant for all bright adaptation contexts and approaching invariance for the dimmer adaptation contexts (Figure 3C,D). Invariance prevailed across most adaptation contexts, arguing that the need for a luminance gain postsynaptic to LMCs generalizes across both fast and slow timescales.

### Gain correction at both fast and slow timescales relies on a single luminance-sensitive input

We next investigated the corrective gain control mechanisms that operate at multiple timescales. L3 neurons provide luminance information and rapidly adjust gain for behavioral response to −100% contrast (4). We now asked if the same source of luminance information at fast timescales simultaneously plays a corrective role at slow timescales. If L3 indeed mediates both fast- and slow-timescale gain correction, a function of the L2 response with limitations at both timescales should capture the behavioral response of flies in which the L3 output is blocked.

To formalize this hypothesis, we modelled the L2 response considering its gain control limitations at fast and slow timescales (Figure 2D, Figure 3A,B). Since L2 takes seconds to adapt (Figure 3A,B), we reasoned that at fast timescales L2 neurons would not encode contrast relative to the current background luminance (i.e. Weber contrast) but relative to their adaptation state modulated by stimulus history. We defined this contrast encoded by L2 as apparent contrast *C_a_*, calculated as the difference between the luminance of the moving edge (*I_edge_*) and the background luminance (*I*), and normalized by the adapting luminance (*I_a_*), which we took to be the mean luminance of the stimulus history (Figure 4A, Methods). The apparent contrast is equal to the Weber contrast only when L2 is adapted to the background luminance of the stimuli (*I* = *I_a_*, *C_a_* = *C*). For example, when the fly encounters a contextually dim stimulus (*I* < *I_a_*), the apparent contrast is lower in magnitude than the Weber contrast (|*C_a_*| < |*C*|) (Figure 4A left). Conversely, when the fly encounters a contextually bright stimulus (*I* > *I_a_*), the apparent contrast is higher in magnitude than the Weber contrast (|*C_a_*| > |*C*|) (Figure 4A right). To test if L2 indeed is more sensitive to apparent contrast than Weber contrast, we measured the response of L2 neurons to randomly changing luminance steps and quantified the response as a function of either of the two contrast definitions. L2 response more strongly correlated with apparent contrast than Weber contrast, especially at shorter step durations (0.5 s to 5 s) that allowed insufficient time for adaptation to each luminance (Figure S3). We thus modelled L2 responses as a function of apparent contrast. At slow timescales, L2 gain depends on absolute luminance (Figure 2D, (14)). We computed this dependence based on previous characterization of fully adapted LMCs (14), thus modelling L2 response as a sigmoidal function of apparent contrast that gets steeper with increasing luminance (Figure 4B).

**Figure 4.**
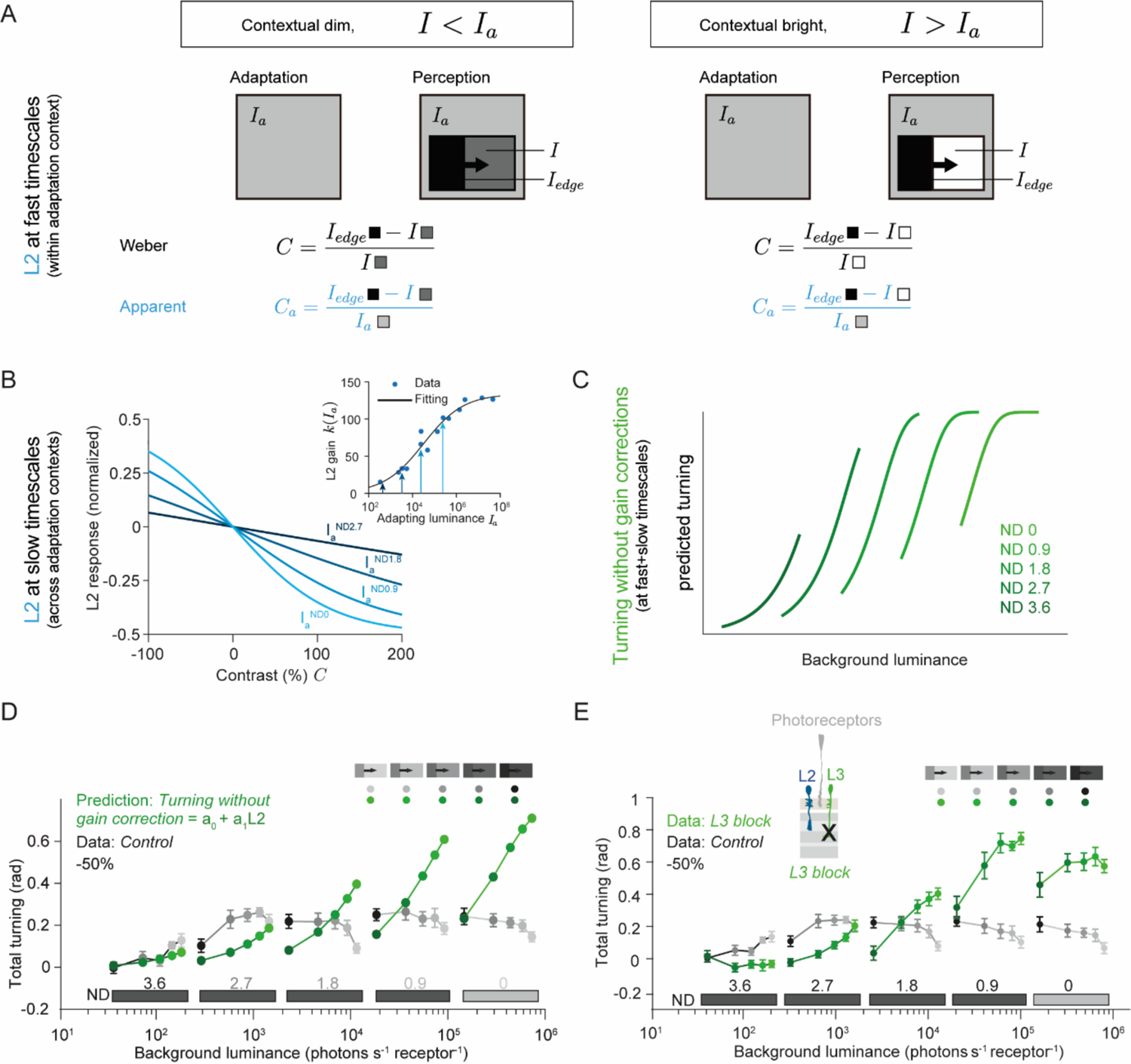
A single neuron type L3 is necessary for gain correction at fast and slow timescales. (A) Definition of apparent contrast (*C_a_*) as the contrast encoded by L2 neurons at fast timescales. As opposed to Weber contrast that normalizes the input difference by the background luminance (*I*), the apparent contrast involves normalization by the luminance to which L2 adapts (*I_a_*). At fast timescales, *I_a_* is different to *I*, and thus the contrasts computed by the two definitions differ. (Left) Contextually dim light, where *I* < *I_a_*. (Right) Contextually bright light, where *I* > *I_a_*. (B) L2 response modeled in four adaptation contexts (ND 0, light blue to ND 2.7, dark blue) as sigmoidal functions of contrast using luminance-dependent gain from the inset. (Inset) Sigmoidal fit (black curve) to L2 gain k(*I_a_*) (the response per unit contrast) as a function of the logarithm of the adapting luminance. Data (blue dots) from (14). The arrows indicate L2 gain values corresponding to the adapting luminances (*I^NDO^_a_* to *I^ND2.7^_a_*) of the four adaptation contexts. (C) Schematic of L2-driven turning behavior in response to a fixed Weber contrast. Turning behavior would vary within and across adaptation contexts, as a result of the L2 gain control limitations at fast and slow timescales. (D) Predicted turning behavior without gain correction (green) modeled as a linear function of predicted L2 responses (B) plotted together with control turning behavior (gray, same data as Figure 3C) in response to −50% contrast in five adaptation contexts. In each context, five different luminance values were tested, presented by the marker shades. The prediction represents the hypothesis that L3 corrects for both fast and slow gain control deficits. (E) Measured total turning response of control and L3-blocked flies to moving OFF edges of −50% contrast, mean ± SEM. Sample size is n=10 flies for each data point.

We then tested if the predicted L2 response with gain control limitations at fast and slow timescales directly translates to turning behavior in the absence of L3 function. If this is true, L2-driven turning behavior without gain correction will vary with luminance within an adaptation context, and the response will also shift up across increasing adaptation contexts (Figure 4C). We then modeled the turning response of flies as a linear function from the predicted L2 response using the −100% behavioral data (from (4), quantified as total turning). Using this model, we then generated the model-predicted turning response of L3-blocked flies to −50% contrast (Figure 4D). The predicted turning responses of L3-blocked flies were lower than the control responses at low absolute luminances (underestimation of contrast) and higher at high absolute luminances (overestimation of contrast), thus showing a bidirectional gain control deficit across luminance (Figure 4D).

To test this hypothesis experimentally, we blocked the output of L3 neurons by acutely expressing *shibire^ts^*, a temperature-sensitive, dominant-negative dynamin allele (23) and measured turning response of these flies to moving edges of −50% contrast. Response of L3-blocked flies did not achieve luminance invariance and scaled with luminance both within and across adaptation contexts (Figure 4E). The turning response of L3-blocked flies was bigger than wildtype turning behavior in bright adaptation contexts, showing a requirement for gain reduction, and smaller in dim adaptation contexts, showing a requirement for gain enhancement (Figure 4E, Figure S4). Overall, the response of L3-blocked flies crossed the control response curve in the intermediate adaptation context. Consequently, L3-blocked response showed a bidirectional deviation from the control response at slow timescales, closely matching the L2-predicted response with gain control limitations at both fast and slow timescales (Figure 4D). Together, this shows that luminance-invariant behavior relies on bidirectional corrective gain control and confirms that L3 is required for both slow and fast gain corrections to achieve luminance-invariant behavior.

### L3 relays information about absolute luminance over several orders of magnitude

Since L3 corrects behavioral response across five orders of magnitude of luminance, and because the absolute luminance values play a key role in determining the direction of the post-receptor gain control, we hypothesized that L3 must have absolute luminance information. We tested this hypothesis by expressing GCaMP6f selectively in L3 neurons and measured calcium signals in response to a broad range of luminances. We exposed flies to 30 s of adapting luminance before showing a −100% contrast OFF step. The dark screen persisted for 3 s before displaying the next adapting luminance. We tested five adapting luminances within one adaptation context, and then repeated the experiment over three more adaptation contexts (Figure 5A). L3 responded with an increase in calcium to the OFF steps (Figure 5B, (22)). We took the calcium response during darkness as the baseline fluorescence (F_0_) across the experiment and quantified the response to different luminances (Figure 5C). The steady-state L3 response gradually decreased with luminance within each luminance regime. Moreover, the response to nearby luminances recorded in a neighboring adaptation context was similar, such that L3 recordings across the entire experiment formed a continuous function of luminance. Only in the brightest adaptation context, the L3 steady-state response curve was somewhat discontinuous, showing a certain degree of gain control (Figure 5C). Overall, this suggests that L3 encodes absolute luminance over three orders of magnitude in its steady-state calcium level, and only transitions to encoding contextual luminance in bright light. Thus, a single neuron, L3, retains luminance information in its baseline calcium signal and is required for a gain correction mechanism shaped predominantly by the absolute luminance values.

**Figure 5:**
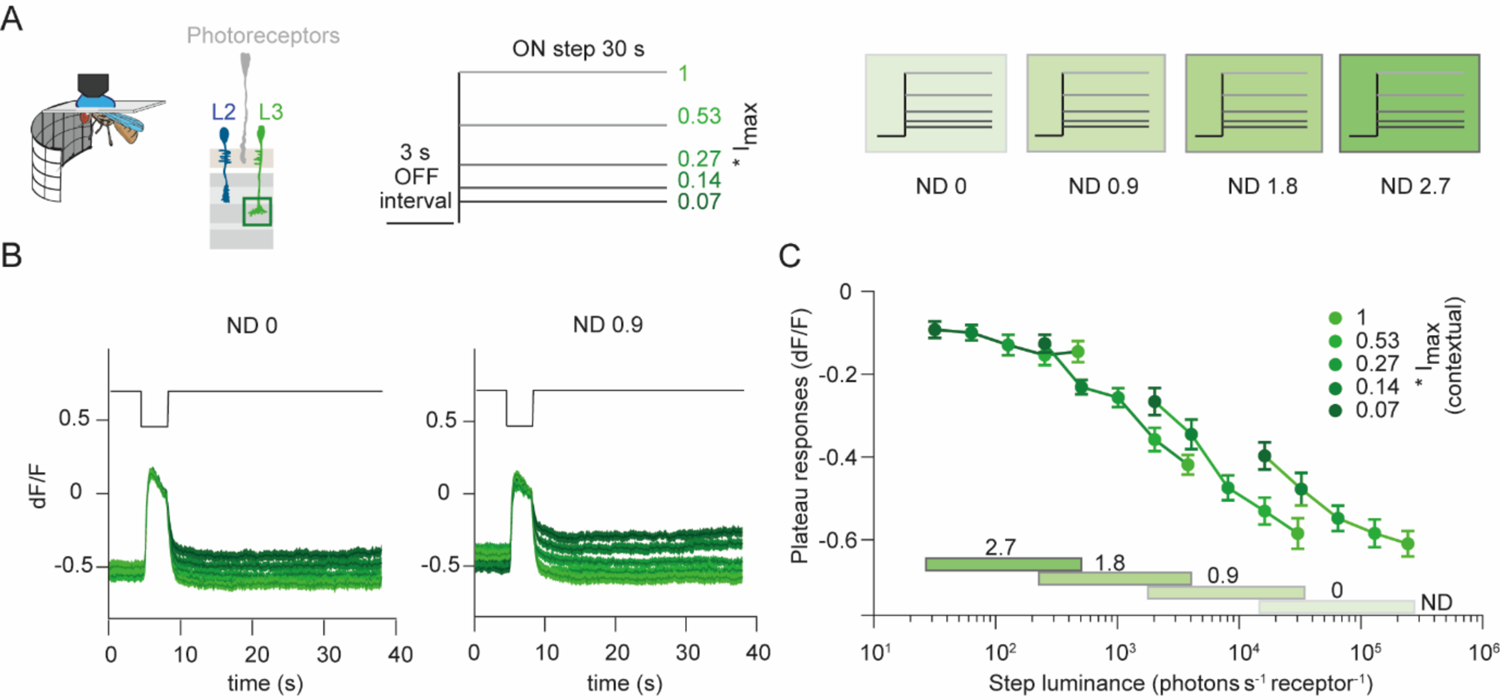
L3 neurons encode absolute luminance. (A) Schematic of the stimuli used for *in vivo* calcium imaging in L3 axon terminals. 30s exposure to one of five different intensity values followed a 3s OFF interval. The procedure was repeated in four different adaptation contexts by using different ND filters. (B) L3 response traces for the five intensity values in the luminance regime ND 0 and ND 0.9. Green shades represent the stimulus intensity within one adaptation context. (C) L3 response quantified in the last second of the 30s luminance exposures for all stimuli over the four regimes. Sample size is n=12 (192), #flies (#ROIs). All traces and plots show mean ± SEM.

### An algorithmic model captures the post-photoreceptor bidirectional gain correction

Our results so far suggest that processing downstream from L3 operates at both fast and slow timescales and implements bidirectional gain control. We found that such bidirectional gain correction across a wide range of luminances was also present at two more contrasts (−80% and −100%) (Figure S5A,B, (4)). However, because behavioral measurement does not allow to measure changes solely at fast timescales, it is still unclear if gain control is bidirectional at both, fast and slow timescales. In order to disentangle the L3 function for fast and slow gain control, we thus extended the L2 model to include the L3 contribution.

In this extended model, we parametrized the ‘L3 contribution’ to behavioral responses. We quantified the L3 contribution by subtracting the total turning of L3-blocked flies from that of control flies (Figure 6A,B). The difference revealed two modes of L3 contribution at −100% contrast (separated by the orange line in Figure 6A). To the right of the orange line, the L3 contribution was higher for dimmer stimuli within each adaptation context. The L3 contribution also gradually shifted down with brighter adaptation contexts (Figure 6A). This suggests that L3 corrected for the response deficits as a function of contextual as well as absolute luminance. To the left of the orange line, the L3 contribution approximated a monotonically increasing function of luminance and did not additionally scale with contextual luminance (Figure 6A). Thus, the L3 contribution appears dichotomous in that L3 provides slow luminance gain control at absolute low luminance, and both fast and slow luminance gain control at high luminance.

**Figure 6.**
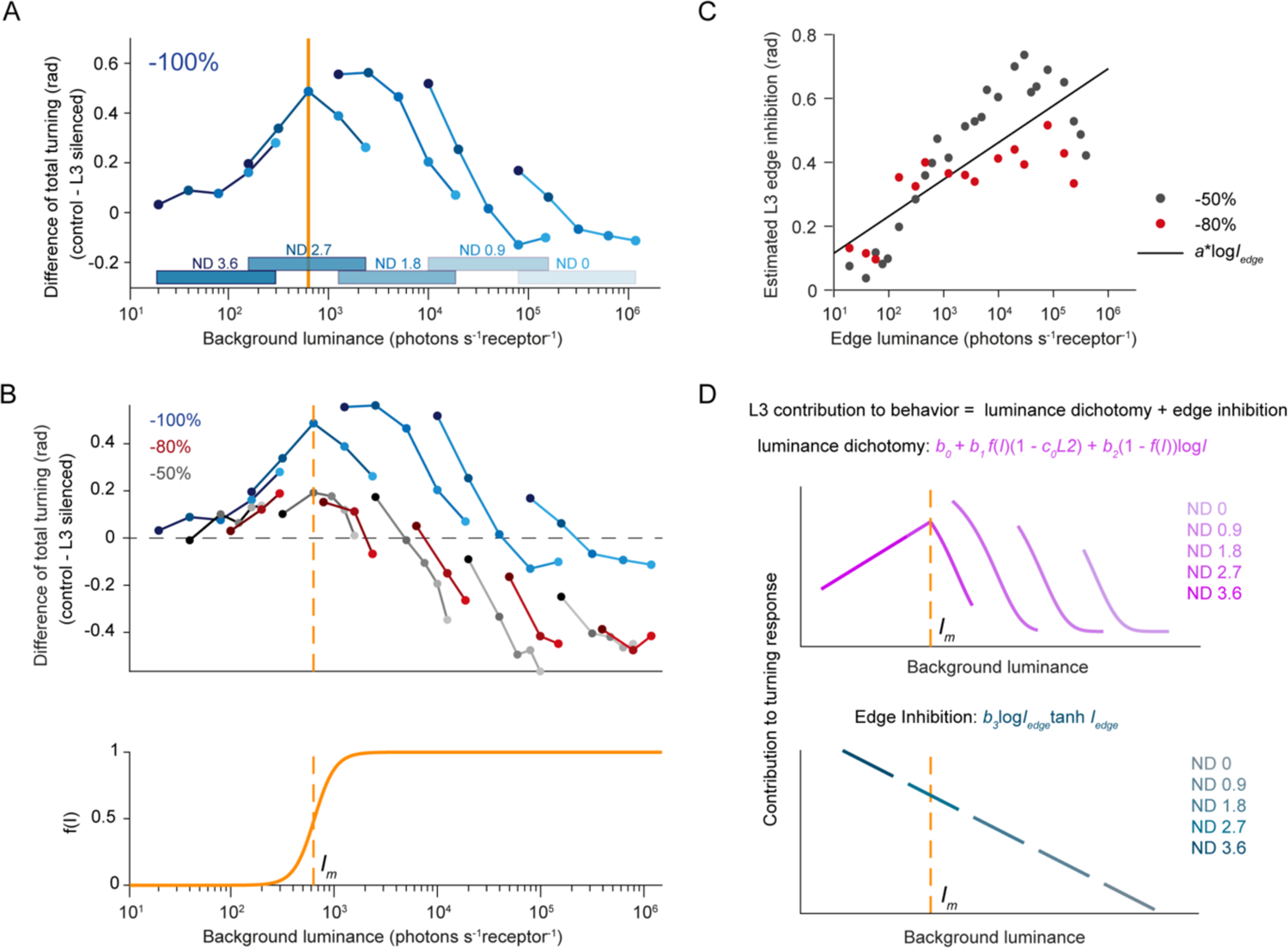
An algorithmic model based on the dichotomous gain corrections of L3. (A) Difference between control response and the response of L3-blocked flies to −100% OFF edge stimuli over five luminance regimes (data from (4), quantified as total turning). The orange line partitions the luminance axis into two parts, which represent the dichotomy of the L3 contribution. (B) (Top) The difference between control response and the response of L3-blocked flies to −50% and −80% OFF edge stimuli over five luminance regimes, compared to −100% OFF edge. The orange line partitions the luminance axis into two parts, the same as in A. (Bottom) A sigmoidal function of luminance centered at the partitioning luminance. (C) The estimated edge inhibition of L3 for −50% (gray) and −80% (red) contrasts. Black line is a linear function of log (edge luminance) fitted to the data points. (D) Schematics describing contributions of the different parts of the algorithmic model to turning behavior. (Top) The luminance-dependent dichotomy in L3 contribution. (Bottom) Inhibition from the moving edge through L3. Lighter line colors indicate brighter adaptation contexts and darker colors indicate dimmer adaptation contexts, as represented by the shaded ND numbers. In C and D, log = log_10_.

We found that the L3 contribution shows a similar dichotomy in adjusting behavioral responses across luminances for −50% and −80% contrast (Figure 6B top). The dichotomy occurred at a similar background luminance I_m_ = 6.3*10^2^ photons/s*receptor for all contrasts and illustrates that distinct types of gain control occur at absolute low and high luminances (Figure 6B). The analysis of the L3 contribution to behavioral responses at all contrasts furthermore revealed that the contribution to turning could implement both gain increments and gain decrements. However, the luminance range where L3 decrements the gain decreases with increasing absolute contrast value (Figure 6B top). For example, comparing to the −80% contrast, the range at which L3 decrements the gain shifted to brighter absolute luminances by ∼1.5 orders of magnitude for −100% contrast. Importantly, edge luminance decreases with increasing contrasts, and is virtually 0 for all −100% stimuli. We therefore postulated that luminance information from the edge itself exerts a negative, or inhibitory L3 contribution that is more prominent in the case of −50% and −80% contrast. We found this inhibitory effect, which we call “edge inhibition”, to be proportional to the logarithm of the edge luminance (Figure 6C, also see Methods).

Building on these observations, we now captured the role of luminance-sensitive pathway with two parallel channels implementing the above-described dichotomy at low and high luminances. We used a single steep sigmoidal function of luminance f(*I*) with its inflection point at the dichotomy luminance *I_m_*(Figure 6B bottom). At low luminances, (1 − *f_I_)) log I* dominates and thus constitutes a channel that implements a luminance-only gain increase independent of contrast. Differently, at high luminances, f(*I*)(1 − b_0_L_2_ dominates and thus constitutes a channel that rescales contrast-driven behavior specific to contextual luminance changes found in dynamic vision, (Figure 6D top). Finally, we linearly combined the contributions from the apparent contrast pathway characterized by a linear function of *L_2_* (Figure 4C, D), the two parallel channels implementing the luminance dichotomy (Figure 6D top) and the inhibition by the moving edge (Figure 6D bottom) to predict the turning behavior.

To determine the scaling constants in the linear combination, we first refitted the linear function of L2 (Figure 4D) on the L3-blocked fly responses using the extended dataset of uniform-contrasts (−100%, −80%, and −50%). Refitting the model for behavioral responses based on L2 responses (Figure 4C,D) on this extended dataset resulted in very similar coefficients (*a_0_* = −0.064, *a_1_* = 0.842 after refitting to −100%, −80%, and −50% contrasts vs *a_0_* = −8.1 × 10^)*^, *a_1_* = 0.781 previously to −100% contrast only). We then fitted the L3 contribution (luminance dichotomy + edge inhibition, Figure 6D) on the difference between the L3-blocked fly responses and the corresponding responses of control flies on the same behavioral datasets. The model captured the predominant underestimation of −100% contrast found in the turning response of L3-blocked flies (Figure 7A, compare to data in Figure S5B) as previously (4). Importantly, the model also captured the bidirectional deviation of turning of L3-blocked flies from turning of controls at −50% and −80% contrasts (Figure 7B,C compared to Figure 4E and Figure S5A). To validate the model prediction on an independent behavioral dataset, we used the experiment where different contrasts were randomly interleaved (Figure 2B), for which we now also measured the turning response of L3-blocked flies (Figure 7D). When we trained the model on the uniform-contrast datasets, it accurately predicted both under- and overestimation of these mixed-contrast stimuli by L3-blocked flies (Figure 7E). Quantitatively, the model captured around 87% of the variance when validated within the three uniform-contrast datasets (R^2^=0.84 to 0.86, depending on contrast; R^2^ for all uniform-contrast datasets together shown in Figure 7F), and around 85% of the variance of the mixed-contrast dataset (R^2^=0.85, Figure 7G). Thus, the model was generalizable to the different stimulus paradigms tested in our experiments. When we trained the model on the mixed-contrast data and validated it on the uniform-contrast data, the model moderately predicted the features of these independent datasets (R^2^ =0.68, Figure S6 A-C), suggesting that the uniform-contrast data comprising luminance invariance at both slow and fast timescales constrained the model better than the mixed-contrast data.

**Figure 7.**
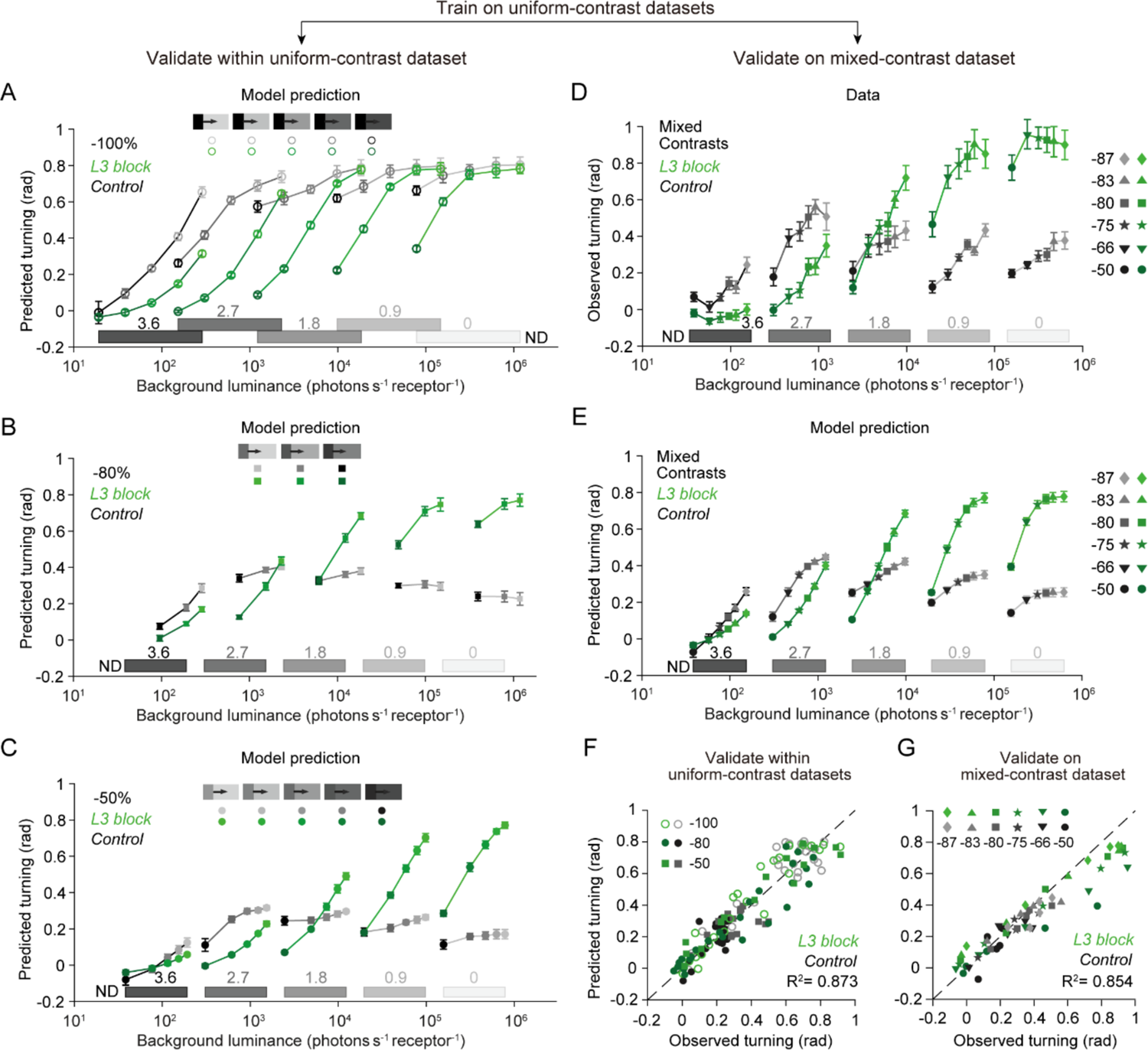
The model captures contrast underestimation and overestimation of L3-blocked flies. (A) Predicted total turning in response to −100% contrast, for control flies (gray) and L3-blocked flies (green). The model was trained on randomly chosen half of the uniform-contrast dataset (−100%, −80%, and −50% together) and predicted the rest half on each iteration. Data are from (4) and shown in Figure S5B. R^2^=0.88%. (B) Same as A, but for −80% contrast. Data are shown in Figure S5A. R^2^=0.84%. (C) Same as A, but for −50% contrast. Data are shown in Figure 4E. R^2^=0.86%. (D) Total turning of control (gray) and L3-blocked (green) flies in response to moving OFF edges of mixed contrasts, measured in five adaptation contexts. Control data same as in Figure 2B. Data are shown as mean ± SEM, sample size n = 10 flies per genotype. (E) Predicted turning response to the mixed contrasts, after training on uniform contrasts, for control flies (gray) and L3-blocked flies (green). (F) Mean predicted turning response from A-C plotted against observed response (from Figure S5 and Figure 4E) to the uniform-contrast stimuli. (G) Mean predicted turning response from E plotted against the corresponding observed response from D. See also Figure S6. The dashed lines in F and G are the unity lines. The predictions in A-C and E are shown as mean ± SD over 30 iterations.

To understand how critically the model’s performance depends on its different components, we performed a sensitivity analysis for all the model’s parameters. The model’s ability to predict individual aspects of the data, such as its ability to predict underestimation and overestimation of contrasts in L3-blocked flies, or the dichotomy, could be attributed to different parameters (see Methods and Table S4, Figure S7).

### The model reveals that L3 implements a bidirectional correction at both fast and slow timescales

Whereas L3 implements a bidirectional gain control (Figure 4D,E, Figure S5), it remains unclear from the experiments alone if the gain control is bidirectional at both fast and slow timescales. We next used the model to decompose the L3 contribution into slow and fast components and asked under which stimulus conditions L3 implements gain increment or gain decrement. We first constructed an ‘average model’ by averaging the scaling coefficients previously obtained over 30 iterations of fitting the model (in Figure 7). Then, we generated model predictions assuming that the contrast-sensitive channel fully adapts, i.e. under the condition *I_a_* = *I* (Figure 8A,B). Under this assumption, the apparent contrast equals Weber contrast (*C_a_* = *C*), and L3 only implements gain corrections at slow timescales. Comparison of the predicted turning response of control (blue) and L3-blocked (orange) flies to −50% contrast revealed that, although the L2 channel was fully adapted under these luminance conditions, the L3-blocked flies would respond less than controls to stimuli in low luminance and respond more to stimuli in high luminance (zoom-in boxes in Figure 8B and Figure S8B,D top). In other words, L3 corrected for the underestimation in low absolute luminance and for the overestimation in high absolute luminance. This corroborated our finding that L3 implements a bidirectional gain control at slow timescales.

**Figure 8.**
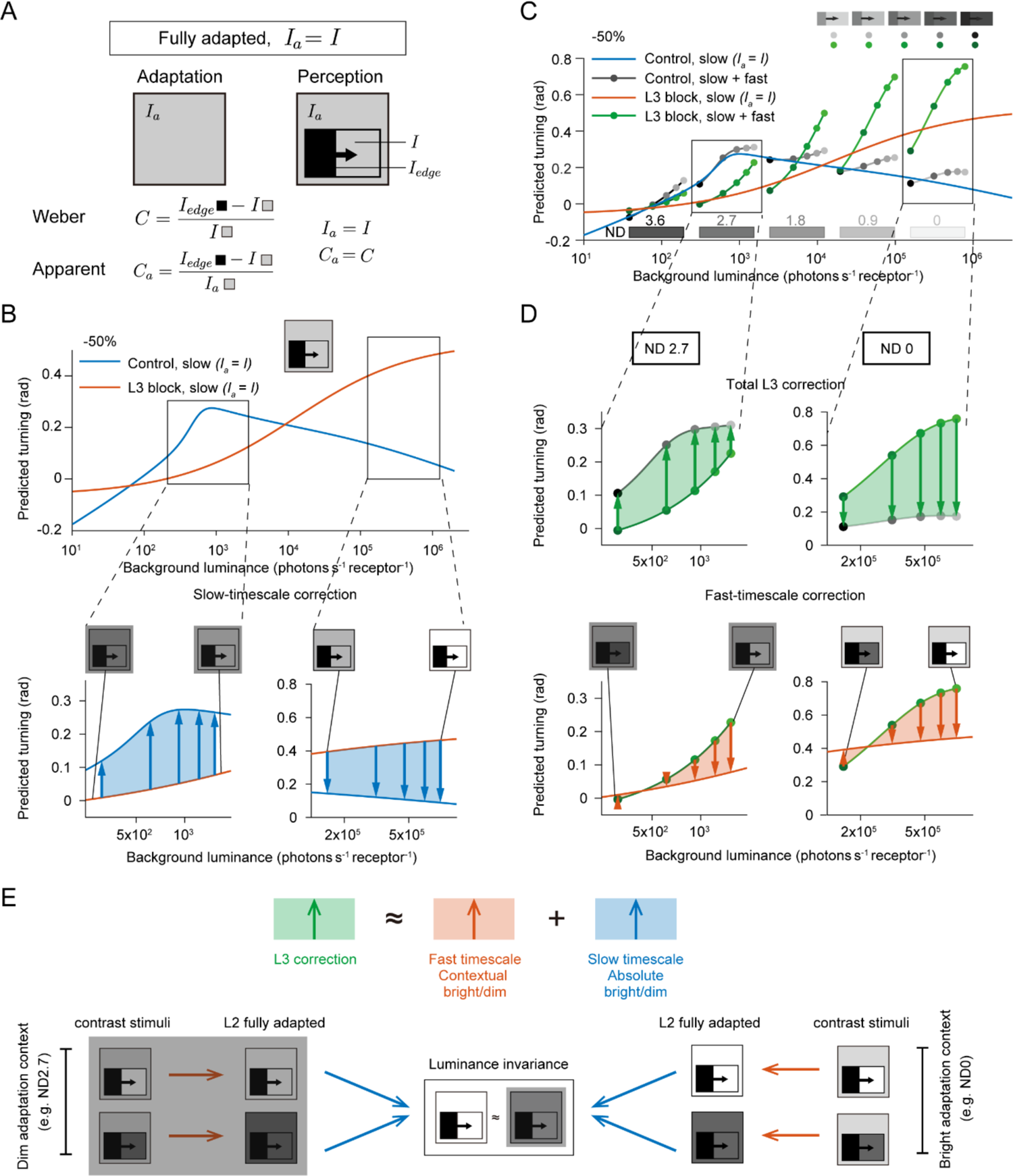
Decomposing the corrective mechanism of L3 into slow-timescale and fast-timescale components. (A) The apparent contrast (*C_a_*) equals to the Weber contrast (*C*) when the contrast-sensitive L2 channel is fully adapted (*I_a_* = *I*). (B) (Top) Predicted turning response to −50% contrast by the average model (see Methods) under the condition of A ((*I_a_* = *I*), for control files (blue) and L3-blocked flies (orange). Boxes indicate zoomed-in intervals in the panel below. (Bottom) Comparison of predicted turning responses of control (blue) and L3-blocked flies (orange) when *I* = *I_a_* shows that the L3-mediated slow-timescale gain correction operates in two directions: gain increments in dim conditions (left, 3.1*10^2^ to 1.5*10^3^ photons s^-1^ receptor^-1^: luminance range similar to ND2.7 in C,D) and gain decrements in bright conditions (right, 1.6*10^5^ to 7.8*10^5^ photons s^-1^ receptor^-1^: luminance range similar to ND0 in C,D). (C) Predicted turning response to −50% contrast by the model at slow + fast timescales, for control files (gray) and L3-blocked files (green), superimposed onto the predictions in B. Boxes indicate zoomed-in intervals in D. (D) (Top) Total L3 correction illustrated as the discrepancy between the predicted control (gray) and L3-blocked fly turning responses at slow + fast timescales (green). (Bottom) Comparison of predicted L3-blocked fly responses between slow + fast timescales (green) and slow timescales enabling complete adaptation (*I* = *I_a_*, orange) shows that the fast-timescale correction operates in two directions: gain increments in contextual dim conditions (the dimmest stimulus in each of the chosen contexts, ND2.7 and ND0) and gain decrements in bright conditions (the four brighter stimuli in each context. In B and D, the arrows point in the direction of gain control: upwards for gain increment, downwards for gain decrements. (E) Schematic summarizing the L3-dependent luminance gain corrections. (Top) The total L3 correction depicted as the sum of the fast-timescale correction and the slow-timescale correction. (Bottom) The fast-timescale correction (orange arrows) ensures invariance between quickly changing background luminances. The slow-timescale correction (blue arrows) ensures invariance among different steady background luminances.

Using the decomposed L3 contribution at slow timescales alone, we next identified the L3 contribution at fast timescales. Specifically, we overlaid the turning response predictions from the average model to −50% uniform contrasts under two conditions: (1) predicted turning response assuming complete adaptation at slow timescales only (*I_a_* = *I*, Figure 8B,C) and (2) predicted turning response at both slow and fast timescales (Figure 8C). The similarity of control fly predictions under the two conditions (blue vs gray, Figure 8C) suggested robust responses across both slow and fast timescales (Figure 8C, Figure S8A-B). On the contrary, the predicted L3-blocked fly response at slow and fast timescales was visibly dissimilar to the predicted response under complete adaptation, and the dissimilarity varied with contextual luminances (orange vs green, Figure 8C, Figure S8A-B), revealing an additional L3 contribution at fast timescales. We reasoned that the contribution of L3 at fast time scales (orange vs green, Figure 8C,D bottom) can be extracted from subtracting the L3 contribution at slow time scales (*I_a_* = *I*), from the total L3 contribution (gray vs green, Figure 8C,D top). This difference represents the discrepancy between the response to apparent contrast and to Weber contrast within an adaptation context. At these fast timescales, L3 corrected for the underestimation of the dimmest stimulus and the overestimation of the brighter stimuli within every adaptation context (orange vs green in Figure 8D, bottom; S8C, middle). Thus, the L3-dependent gain correction is bidirectional at both fast and slow timescales, such that the gain is increased for stimuli dimmer than the adapting luminance and decreased for those brighter than the adapting luminance.

Taken together, the L3-mediated bidirectional gain correction plays distinct roles at both fast and slow timescales: the fast-timescale gain correction achieves encoding of Weber contrast at quickly changing luminances, and the slow-timescale correction reduces the dependence of luminance gain on the absolute adaptation state. By combining these two corrections, the model reproduces the luminance-invariant fly behavior (Figure 8E).

In sum, our results uncover a post-photoreceptor luminance gain control mechanism that confers robustness to visual processing in dynamic conditions, encompassing both fast and slow timescales. Our work demonstrates that fly behavior achieves luminance invariance and better detection of dim contrast stimuli than expected solely from contrast-sensitive (L2) physiology, owing to the multifaceted gain corrections across luminances. The luminance-sensitive L3 neurons provide signals necessary for both gain increments and decrements at all timescales.

## Discussion

In the present study, we have demonstrated that luminance gain control in the circuitry past photoreceptors is crucial for stable processing of visual contrasts over a broad luminance space, ensuring luminance-invariant behavioral responses. Luminance-invariant vision is challenged at both slow (>30s) and fast (<1s) timescales, and in two seemingly opposite ways: contrasts are (1) underestimated in dim light, requiring a corrective gain increment and (2) overestimated in bright light, requiring a corrective gain decrement. We have shown that a single luminance-sensitive L3 channel provides signals for these diverse gain-corrective operations. An algorithmic model implementing these diverse operations proposes that the luminance signals play distinct roles: a nonlinear interaction with contrast-sensitive signals implements the gain correction at fast timescales, whereas the luminance information alone implements the slow gain correction. Together, our findings show that luminance gain control is a multifaceted computation and is indispensable to stable behavior in changing environments.

### Contrast computation in first-order interneurons is insufficient for luminance invariance at slow and fast timescales

Luminance adaptation starts at the entry point of the visual systems, in photoreceptors. The adaptation processes begin within milliseconds and continue for several seconds or even minutes (3, 13, 16, 24–26). But at what adaptation timescale do photoreceptors produce sufficient gain control for luminance-invariant behavior? Both the comparison of L2 physiology with behavior and the behavior of flies lacking the source of post-photoreceptor luminance information, L3, reveal that photoreceptors do not achieve this goal even after long adaptation times. A post-photoreceptor gain control is indispensable even at slow timescales. At fast timescales (<1s), when photoreceptors are not fully adapted, the gain control past the photoreceptors contributes additionally. Many behaviors act on such fast timescales. For example, behavioral responses to visual stimuli occur within tens of milliseconds, and thus rely on a gain control operation that acts rapidly. Thus, gain control processes at different temporal scales extend beyond the peripheral visual circuitry.

Does peripheral gain control suffice for luminance invariance in any luminance conditions? In big flies, LMC responses stabilizes at high luminance (14), which might either indicate saturation or show that LMC responses eventually becomes luminance invariant. Since we explored a luminance range resembling twilight, most relevant for *Drosophila*’s crepuscular lifestyle, it remains untested if *Drosophila* L2 responses can become luminance invariant at higher luminance. Notably, vertebrate photoreceptors appear to achieve ideal gain control across a wide luminance range, as implied by their threshold-vs-intensity curves following Weber’s law (7, 27). In the “Weber regions” of these curves, the detection threshold varies proportionally with the background intensity. However, in low luminance regions, the same curves rise less steeply than expected from Weber’s law. For example, in the “Rose-DeVries region”, the detection threshold varies with the square root of background luminance (7, 28, 29). Although ideal gain control is also missing at the level of human perception in this luminance region, post-photoreceptor gain control mechanisms might be contributing to improved contrast perception in these dim conditions.

Our work examines fly behavior and physiology of the first-order interneurons, the LMCs L2 and L3, using very similar stimulus parameters and hence offers a direct comparison between the two measures, demonstrating that behavior achieves better luminance invariance than the peripheral gain control. Given the shared presence of luminance ranges challenging invariance in different animals, involvement of additional of gain control downstream of the sensory periphery can be a shared characteristic of their visual systems.

### General benefits of post-photoreceptor luminance gain control

Implementing gain control mechanisms postsynaptic to photoreceptors has multiple benefits. First, besides achieving luminance-invariant contrast processing, gain regulation is necessary to match the input range with a neuron’s operating range. For example, photoreceptors encode light inputs from natural scenes comprising approximately ten orders of intensity variations, with just two orders of voltage response range (5, 30, 31). At advanced processing stages, the input from upstream neurons may again exceed the operating range and subsequent layers of gain control could prevent information loss stemming from this mismatch. Secondly, photoreceptors are local units that respond to single points in visual space. In contrast, post-receptor gain control is thought to operate on a relatively noise-free, spatially pooled signal making it more reliable especially at fast timescales (7). The contribution of post-receptor gain control to luminance-invariant behavior in the vertebrate visual system still remains unexplored. Interestingly however, in line with our findings in the fly visual system, the major gain control in the vertebrate retina is also known to occur in the post-photoreceptor sites such as RGCs, instead of photoreceptors or bipolar cells, especially in low light that renders photoreceptor signals noisy and thus unreliable (32). To increase the signal-to-noise ratio, visual circuitry in both vertebrate retina and invertebrates could spatially pool luminance information in wide field neurons downstream of photoreceptors, such as the horizontal or amacrine cells in vertebrates, or the amacrine-like Dm (distal medulla) or Pm (proximal medulla) neurons in Drosophila (33–36).

### Multifaceted gain control is implemented downstream of a single luminance-sensitive neuron L3

Modeling allowed us to algorithmically link the physiology of the first-order visual interneurons to behavior and to abstract the LMC activity into contrast and luminance information. The model decomposed the contribution of L3 to fast and slow luminance gain controls, a distinction that remained elusive from the experiments alone. The fast-timescale correction achieves a luminance-invariant response within each adaptation context, akin to the encoding of Weber contrast, as opposed to the luminance dependence in the first-order interneurons. On the contrary, the slow-timescale gain correction achieves a luminance-invariant response across adaptation contexts that span at least five orders of luminance. The model assumes an absence of adaptation when correcting gain at fast timescales and complete LMC adaptation to the mean luminance at slow timescales. The combined gain corrections in these two extreme cases can explain the overall L3-mediated correction in our experiments comprising multiple timescales (Figure 8E). Thus, due to its modular composition, we argue that the model can capture for luminance-invariant behavioural responses to diverse temporal stimulation patterns, including those present in natural environments of behaving animals.

In addition to acting at different timescales, gain control increases or decreases neuronal response to the same input, depending on whether viewing conditions turn dimmer or brighter. Both computations take place downstream of the luminance-sensitive L3 neurons. This could be easily explained if feedback (or cell intrinsic) mechanisms were involved. For instance, gain control stemming from neuronal feedback or cell-intrinsic adaptation processes arises due to deviations from a target activity or output (e.g., photoreceptor-horizontal cell feedback loop) (37–39), and such feedback automatically goes up or down when the output itself goes up or down. However, L3 neurons themselves rely little on circuit interactions such as feedback (40). Thus, the bidirectional gain control downstream of L3 is potentially a feed-forward circuit mechanism. In that case, separate excitatory and inhibitory channels downstream of L3 would implement the two oppositely signed gain corrections. A split into separate channels downstream of L3 is also beneficial when the need for different types of gain control is not symmetrical, that is when the post-photoreceptor gain correction is differentially required to increase and decrease luminance gain. For example, dark adaptation in photoreceptors involves slow recovery of bleached visual pigments, and thus takes much longer (up to an hour) than light adaptation (3, 41, 42). In this scenario, the post-photoreceptor gain control may play a more extensive role in dark adaptation at fast timescales than in light adaptation.

Post-receptor gain control in vertebrates also involves non-intrinsic, circuit components. For example, gain control in bipolar cells includes a potential input from horizontal cells (43). Gain control in retinal ganglion cells (RGCs) relies on upstream neurons that form the summation pool for RGCs (44). It will be interesting to test if and how these circuit motifs implement bidirectional gain control. How distinct gain-corrective mechanisms are combined to ultimately guide luminance-invariant behavior remains to be explored and will be facilitated by *Drosophila* genetics and the rapidly advancing connectomics. As one candidate, Tm9 receives the main input from L3 and also exhibits a sustained response (40, 45). It could scale up contrast information coming indirectly from L1, which also contributes to the Tm9 response (45), and therefore could achieve gain increments necessary to establish invariance in contextual dim light and to improve dim light vision. Interestingly, different types of wide-field Dm neurons innervate medulla layer M3 (33), which houses the L3 axon terminals and Tm9 dendrites. Thus, Dm neurons could pool luminance information and pass it to Tm9 neurons, which have been shown to have wide receptive field properties (45, 46). This is also consistent with the idea that the L3-mediated gain control benefits from spatial summation as discussed earlier.

L3 response is a monotonic function of luminance, thus lacking any intrinsic dichotomy. Yet, the gain control implemented by L3 abruptly changes at a specific luminance *I_m_*, from being strictly incremental and absolute luminance-dependent at low luminances *I* < *I_m_* to being bidirectional and contextual luminance-dependent at high luminances *I* > *I_m_*. The computations at luminances *I* < *I_m_* might thus heavily be driven by a low luminance-dependent recruitment of wide-field neurons, such as the Dm neurons. Using larger receptive fields at low luminance helps the system reduce noise and achieve better sensitivity while trading off spatial resolution. Such a mechanism would also be in line with the observation that the extent of spatial summation in the visual systems increases as the luminance decreases (47, 48).

### L3 encodes absolute luminance and improves dim light vision

Flies did not achieve luminance invariance in behavior in dim adaptation contexts, reminiscent of the Rose-de Vries region in human perception. However, the L3-mediated gain increment in these contexts still helped improve detection of very dim stimuli. Thus, the L3 pathway could be a strategy of the invertebrate visual system to improve scotopic vision while relying on a single photoreceptor system that covers a wide range of luminances, as opposed to the vertebrate rod-cone systems. Besides, spatial summation predicted to occur downstream of L3 can further enhance dim light detection, by improving signal-to-noise ratio. In line with this, nocturnal hawkmoth vision works better than the photoreceptors would allow, due to spatial summation (49). Importantly, L3 encodes absolute luminance over several orders of luminance. Absolute luminance sensitivity is also found in vertebrate retina e.g. in intrinsically photosensitive retinal ganglion cells (ipRGCs). However, different ipRGCs from the population tile the luminance space so that they can collectively encode the broad luminance range with sufficient resolution (50). In contrast, L3 encodes nearly three orders of luminance with its limited operating range, arguing that L3 is either remarkably precise or that the fly visual system compromises on the ability to discriminate between close-by luminances.

Together, our study uncovers a gain control strategy that ensures both better stimulus detection at low luminances and luminance invariance in a broad luminance range. Together, this leads to a better behavioral performance than can be done by photoreceptor gain control alone. Most importantly, the mechanism ensures stable performance both when the day slowly advances, as well as when an animal actively explores the environment, thus being crucial across vastly different natural timescales.

## Materials and Methods

### Experimental model

Flies were raised at 25°C and 65 % humidity on standard molasses-based fly food in climate chambers with a 12h:12h light-dark cycle. Two-photon experiments were conducted at room temperature (20°C) and behavioral experiments at 34°C. Female flies 2-4 days after eclosion were used for all experimental purposes. Fly lines used for genetic silencing were *L3^0595^-Gal4,* described in (22), and *UAS-shi[ts]* from BDSC (#44222). For imaging experiments, GCaMP6f (BDSC #42747) was expressed using *L2^21Dhh^-Gal4* (51) and *L3^MH56^-Gal4* (52). Detailed genotypes are given in Table 1.

**Table 1:** Genotypes used in this study.

| Name | Genotype | Figure |
| --- | --- | --- |
| **Imaging** |  |  |
| *L2>>GCaMP6f* | *w+; UAS-GCaMP6f / +; L2<sup>21Dhh</sup>-Gal4 / +* | Fig 2, 3, S1, S2, S3 |
| *L3>>GCaMP6f* | *w+; L3<sup>MH56</sup>-Gal4 / +; UAS-GCaMP6f / +* | Fig5 |
| **Behavior** |  |  |
| UAS-shibire<sup>ts</sup> control | *w+; + / +; UAS-shi<sup>ts</sup> / +* | Fig 2, 3, 4, 6, 7, S1, S4, S5, S6, S7 |
| L3-Gal4 control | *w+; + / +; L3<sup>0595</sup>-Gal4 / +* | Fig S4 |
| L3 silencing | *w+; + / +; L3<sup>0595</sup>-Gal4 / UAS- shi<sup>ts</sup>* | Fig 4,6,7, S4, S5, S6, S7 |

### Behavioral experiments

Behavioral experiments were performed as described in (4). In brief, all experiments were conducted at 34°C, a restrictive temperature for *shibire^ts^* (23). Female flies were cold anesthetized and glued to the tip of a needle at their thorax using UV-hardened Norland optical adhesive. A 3D micromanipulator positioned the fly above an air-cushioned polyurethane ball (Kugel-Winnie, Bamberg, Germany), 6 mm in diameter, and located at the center of a cylindrical LED arena that spanned 192° in azimuth and 80° in elevation (53). The LED arena (IO Rodeo, CA, USA) consisted of 570 nm LED panels and was enclosed in a dark chamber. The pixel resolution was ∼2° at the fly’s elevation. Rotation of the ball was sampled at 120 Hz with two wireless optical sensors (Logitech Anywhere MX 1, Lausanne, Switzerland), positioned toward the center of the ball and at 90° to each other (setup described in (54)). Custom written C# code was used to acquire ball movement data. MATLAB (Mathworks, MA, USA) was used to coordinate stimulus presentation and data acquisition. Data for each stimulus sequence were acquired for 15-20 minutes, depending on the number of distinct epochs in the sequence (see ‘visual stimulation’ for details).

### Visual stimulation for behavior

The stimulation panels consist of green LEDs that can show 16 different, linearly spaced intensities. The presented luminance was measured as previously described (4). Briefly, candela/m^2^ values were first measured from the position of the fly using a LS-100 luminance meter (Konika Minolta, NJ, USA). Then, these values were transformed to photons incident per photoreceptor per second, following the procedure described by (55) and using parameters specific to the *D. melanogaster* compound eye (56). The highest native LED luminance was approximately 11.77 * 10^5^ photons * s^-1^ * photoreceptor^-1^ (corresponding to a measured luminance of 51.34 cd/m2), and the luminance meter read 0 candela/ m^2^ when all LEDs were off. Neutral density filter foils (Lee filters) were placed in front of the panels to attenuate luminance. The foils were used individually or in combinations to achieve optical densities of 0.9, 1.8, 2.7 and 3.6, thus creating luminance regimes that spanned about four orders of magnitude.

Fly behavior was measured in an open-loop paradigm where OFF edges varying in either contrast or luminance, depending on the experiment, were presented in a randomized order. For every set of OFF edges, each edge was presented for around 60 to 80 trials. Each trial consisted of an initial static pattern (i.e. the first frame of the upcoming pattern) shown for 500 ms as background, followed by 750 ms of edge motion. Inter-trial intervals were 1s long and always consisted of a dark screen. All edges from a set were presented in a mirror-symmetric fashion (moving to the right, or to the left) to account for potential biases in individual flies or introduced when positioning on the ball.

1. Uniform-contrast edges: These were two sets comprising −50% and −80% (Weber) contrast edges separately. In the −50% set, the edges were made of 5 different luminance values (i.e. five unique epochs), moving onto backgrounds twice as bright as the respective edges. The edges assumed 7%, 14%, 20%, 27% and 34% of the highest LED intensity (corresponding to the luminances: 0.98, 1.96, 2.94, 3.92 or 4.90 *10^4^ photons*s^-1^*receptor^-1^ luminance). The −80% set comprised three unique edges with luminances 0.98, 1.96 or 2.94 *10^4^ photons*s^-1^*receptor^-1^, which moved onto backgrounds that were five times as bright as the corresponding edges. The data for behavioral response to −100% Weber contrast were published previously (4), but in this study, total turning was quantified and plotted as opposed to peak turning in (4).
2. Mixed-contrast edges: The set comprised of six distinct epochs, each with a different Weber contrast value (50%, 66%, 75%, 80%, 83% and 87%). Here, the edge luminance was maintained constant at 7% of the highest LED intensity, across epochs, and the background luminance varied.

### Behavioral data analysis

Fly turning behavior was defined as yaw velocities that were derived as described in (54), leading to a positive turn when flies turned in the direction of the stimulation and to a negative turn in the opposite case. Turning elicited by the same epoch moving either to the right or to the left were aggregated to compute the mean response of the fly to that epoch. Time traces of turning responses are presented as angular velocities (rad/s) averaged across flies ± SEM.

Total turning was calculated between 200 ms post the edge motion onset (to account for response latency) and the time point when the turning speed averaged over all trials drops to 0, restricted to a 1 and 1.5s window after the edge motion onset, that the quantification remained unaffected by spurious drops of velocity occurring in noisy traces. Quantification of total turning offered a relatively noise-free metric, considering flies did not make much net directional movement in some trials. In specific cases where statistical tests were performed, total turning was first tested for normal distribution using a Kolmogorov-Smirnov test. To test if the normally distributed turning response varied with luminance within an experimental condition, Welch’s ANOVA was performed (Figure 3C,D), followed by Bonferroni-Holm correction to account for multiple tests (5 tests in each plot, corresponding to 5 adaptation contexts). Flies with a forward walking speed less than 3 mm/s computed in the last 200 ms of the inter-trial interval were discarded from the analysis. This resulted in rejection of approximately 15% of all flies, independent of genotype.

### Two-photon imaging

Female flies were anesthetized on ice before placing them onto a sheet of stainless-steel foil bearing a hole that fit the thorax and head of the flies. Flies were head fixated using UV-sensitive glue (Bondic). The head of the fly was tilted downward, looking toward the stimulation screen and the back of the head was exposed to the microscope objective. To optically access the optic lobe, a small window was cut in the cuticle on the back of the head using a fine razor blade and sharp forceps. During imaging, the brain was perfused with a carboxygenated saline-sugar solution composed of 103 mM NaCl, 3 mM KCl, 5 mM TES, 1 mM NaH2PO4, 4 mM MgCl2, 1.5 mM CaCl2,10 mM trehalose, 10 mM glucose, 7 mM sucrose, and 26 mM NaHCO3. For dissections, the same solution lacking calcium and sugars was used. The pH of the saline equilibrated near 7.3 when bubbled with 95% O2 / 5% CO2. The two-photon experiments were performed using a Bruker Investigator microscope (Bruker, Madison, WI, USA), equipped with a 25x/NA1.1 objective (Nikon, Minato, Japan). An excitation laser (Spectraphysics Insight DS+) tuned to 920 nm was used to excite GCaMP6f, applying 5-15 mW of power at the sample. Emitted light was sent through a SP680 shortpass filter, a 560 lpxr dichroic filter and a 525/70 emission filter. Data was acquired using PrairieView software at a frame rate of 8 to 12 Hz.

### Visual stimulation for imaging

For comparability between behavior and physiology, stimuli for imaging experiments were shown using a cylindrical LED panels arena with the same dimensions as the one used for behavior. The arena (IO Rodeo, CA, USA) consisted of 470 nm LEDs with pixel resolution of ∼2° at its center. The LEDs can show 16 different, linearly spaced intensities. The arena was positioned below the stage of the two-photon microscope, such that the fly position under the two-photon objective was approximately at the same height as the upper border of the arena Two layers of blue filter (Rosko, Indigo no. 59) and a 0.6 neutral density (ND) filter foil (Lee filters) were placed in front of the arena to minimize bleedthrough onto the detectors, such that the maximum luminance (highest LED level in the regime ND 0) was 2.43 * 10^5^ photons * s^-1^ * photoreceptor^-1^. Further neutral density filters were added to achieve dimmer luminance regimes. MATLAB coupled with a data acquisition device (National Instruments, NI USB-6211) coordinated stimulus presentation, stimulus data acquisition and a start trigger to the microscope software to synchronize imaging with stimulus presentation.

All stimuli were full-field flashes. For L2 imaging with adapting conditions (Figures 2, 3, S1, S2), a set of contrast steps was shown at five adaptation contexts ND 0 (the brightest), ND 0.6, ND 1.2, ND 1.8 and ND 2.4 (the dimmest). In each context, 30 s long exposure to 53% of the maximum luminance was alternated with 30 s long ON or OFF of contrast steps, where the contrast value was one of −100%, −87.5%, −75%, −62.5%, −50%, −25%, 25% and 50% Weber contrasts, randomly chosen. Each step was shown once, thus imaging time per adaptation context was ∼8 minutes. Sequence of the adaptation contexts was also randomized for each fly.

For L2 imaging with dynamic conditions (Figure S3), a ‘random luminance steps’ stimulus was shown in two adaptation contexts, ND 0.9 and ND 1.5. In each context, pseudo-randomized stimulus sequences comprising between 45-72 steps of six unique luminance values (0%, 7%, 14%, 27%, 53% and 100% of the luminance of the highest LED level) were tested with four different step durations: 0.5s, 1s, 5s and 10s. In each recording, a 10s dark interval preceded the sequence. The imaging time per recording varied between 36s and 8 minutes, based on the chosen step duration.

For L3 imaging, the stimulus was made of a dark 3s interval of ∼zero luminance alternating with five luminances, each shown for 30 s. The luminances were 7%, 14%, 27%, 53% or 100% of the luminance of the highest LED level in each of the four luminance regimes ND 0, ND 0.9, ND 1.8 and ND 2.7. Sequence of the regimes as well as sequence of the luminances in each regime was randomized for each fly. The five luminances were shown three times each.

### Imaging data analysis

Two-photon data were analyzed using custom-written Matlab code. Images were corrected for fly movements using cross-correlation upon Fourier transformation and aligned to a reference image that was maximum intensity projection of the first 30 images. Regions of interest (ROIs) were selected manually from an averaged aligned image such that each ROI enclosed an axon terminal. Average fluorescence intensity of each ROI was calculated for each frame to acquire fluorescence time traces. Fluorescence time trace of a background region was subtracted from the time trace of each ROI. Such background-subtracted signals and stimulus traces were then interpolated to 10 Hz and trial averaged. Normalized signal was calculated for each ROI as (dF/F) = (F – F_0_)/ F_0_, where F is the fluorescence at each time point and F_0_ was defined separately for each experiment as described below.

<u>L2 imaging:</u> (1) Adapting conditions (Figures 2, 3, S1, S2): F_0_ equaled mean fluorescence of the entire time series of the ROI. (2) Dynamic conditions (Figure S3): Mean fluorescence intensity of the ROI during the dark epochs preceding the random sequence was used as F_0._

<u>L3 imaging:</u> Mean fluorescence intensity of the ROI during the dark epochs was used as F_0._

For both L2 and L3, single ROI signals having negative correlation with stimulus were chosen for further analysis. A minority of ROIs were positively correlated with stimulus, as described previously (45). These ROIs have receptive field outside the stimulation area and were thus discarded. Responses of the chosen ROIs were averaged for each fly, and the fly means were further averaged to obtain mean response traces, shown in Figures 2 and 5 as mean ± SEM. Responses of L2 in dynamic conditions were averaged directly over ROIs to obtain mean response traces (Figure S3A). Furthermore, the correlation of L2 step responses in dynamic conditions with stimulus contrasts was calculated as Pearson’s correlation coefficient (R), to be compared across two contrast definitions: apparent and Weber contrast (Figure 4A, S3B,C). The R values were first tested for normal distribution using a Kolmogorov-Smirnov test. To test if the normally distributed R differed between the two contrast definitions within an experimental condition, paired-sample Student t-tests were performed (Figure S3B,C), followed by Bonferroni-Holm correction to account for multiple tests (a total of 8 paired-sample tests). To test if the correlation of L2 responses with Weber contrasts differed between the experiments with different step durations, two-sample Student t-tests were performed, followed by Bonferroni-Holm correction to account for multiple tests (a total of 12 two-sample tests).

### The luminance-contrast integration model

1. Modeling L2 fly responses The L2 neuron computes contrast relative to the adapting luminance, which was assumed to be the mean luminance of the adaptation context being tested during an experiment. The adapting luminance *I_a_* was thus calculated as the temporal average of different luminance values shown within an adaptation context:

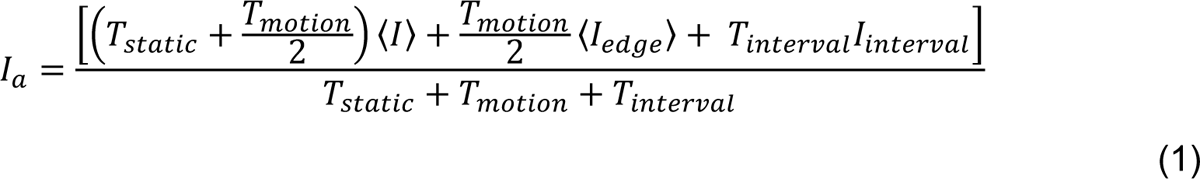

where the static background duration was *T_static_* = 0.5 s, the edge motion duration was *T_motion_* = 0.75 s, the inter-trial interval *T_interval_* = 1 s, and the luminance of the inter-trial *I_interval_* = 0. 〈*I_edge_*〉 was the mean luminance of OFF edges and 〈*I*〉 the mean luminance of the backgrounds specific to the experiments. The apparent contrast was then calculated as:

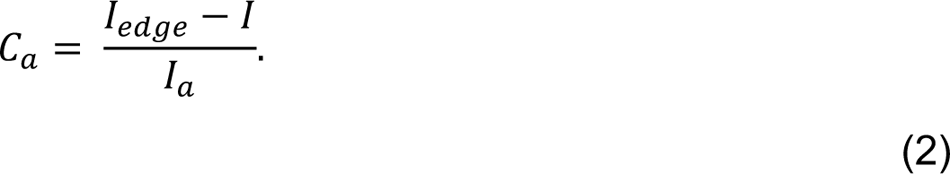 This is different to the Weber contrast, defined as

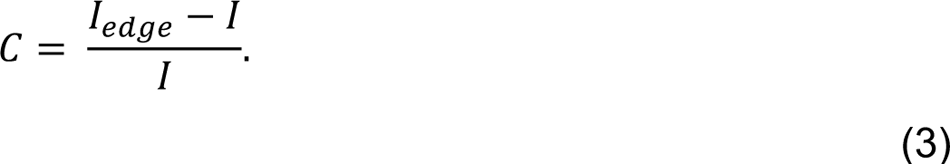 In Figures 4, 7 and 8, we modeled the L2 response as sigmoidal functions of contrast (14), which in our case were rescaled to take the form of hyperbolic tangent functions, such that the output spans [-1, 1]:

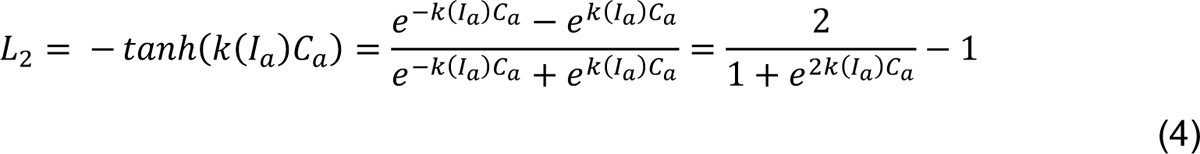

where the gain *k*(*I_a_*) depends on the adapting luminance as measured in previous experimental measurements (14). The gain was fitted on the contrast response curve published in (14) using the equation

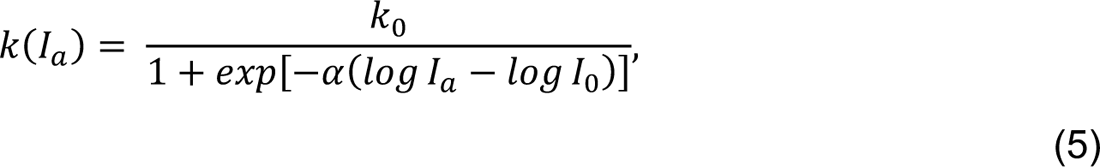

with parameters *k_0_* = 133.9, α = 1.095, and *log I*_0_ = 4.505 (log with base 10, also the same in following paragraphs). Given the difference in species and other differences between our experiments and those of (14), we allowed the gain of L2 to be different from (14) by considering *k_0_* as an additional free parameter during the fitting procedure (see Supplementary Tables).
2. Modeling L3-blocked fly responses to −100% contrasts To generate the predictions in Figure 4D, we fitted the linear function *a_0_* + *a_1_L_2_* on the L3-blocked turning response to −100% contrasts, where *L_2_* denotes the contrast-sensitive LMC output (see Eq. (4)). The luminance-dependent gain of L2 was given by Eq. (5) above. Parameter estimates obtained by fitting are shown in the Table S1. The fitted linear functions were then used to predict the turning response of L3-blocked flies to −50% contrasts in Figure 4D.
3. Modeling the dichotomous L3 contribution to fly responses The dichotomy in Figure 6 was characterized by a steep sigmoidal function *f(I)* (Figure 6B bottom). We determined the midpoint of f(*I*) by visual observation of where L3 contribution to turning behavior switches the mode of gain correction (Figure 6A). The midpoint, or the dichotomy point *I_m_*, was thus defined as the luminance with the logarithm of 2.8 (log *I_m_* = 2.8). To mimic the conspicuous switch at the dichotomy point *I_m_*, we assumed that f(*I*) is very steep (a slope of 10) such that it splits the luminance range into two pronounced parts. The f(*I*) was thus defined as:

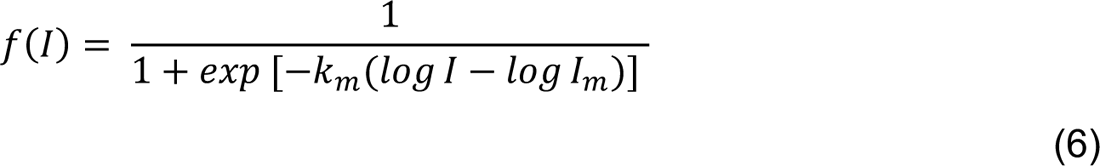

with log *I_m_* = 2.8 and *k*_%_ = 10 (Figure 6B bottom, *I* is expressed in photons receptor^-1^ s^-1^). Then the dichotomous contribution of L3 consisted of two parts partitioned by *f(I): b_1_f(I)(1 − c_0_L_2_*) was the dominant term at high luminances, whereas *b_2_(1 − f(I)) log I* was dominant in dim light.
4. Estimating the edge inhibition of the L3 contribution to fly responses To estimate the effect of edge inhibition, the two parts of the dichotomous L3 contribution were linearly combined allowing also for a constant offset *b_0_*:

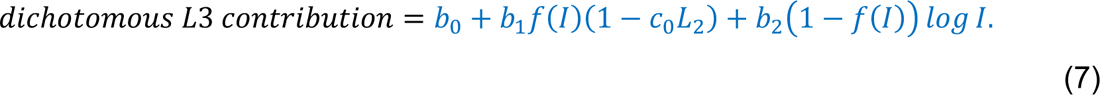 We fitted this model on the measured L3 contribution (control turning minus L3-blocked turning) in response to −100% contrasts (data reused from (4) and analyzed to acquire total turning) and predicted the L3 contribution in −50% and −80% contrast datasets. We then calculated the discrepancy in L3 contribution (−50% and −80% contrasts) between the data (Figure 6B) and the model predictions from Eq. (7). We found that the discrepancy, as an estimate of the edge inhibition effect, is a linear function of the logarithm of the edge luminance, i.e., *a* * log *I_edge_* (R^2^ = 0.603, p = 3.96 × 10^−9^, Figure 6C).
5. Model trained on the uniform-contrast datasets Since the predicted turning without gain control, modeled as linear function of predicted L2 responses (section2), captured the L3-blocked fly turning in response to −50% contrasts (Figure 4D-E), we modeled the L3-blocked turning response using the same linear function of L2 responses:

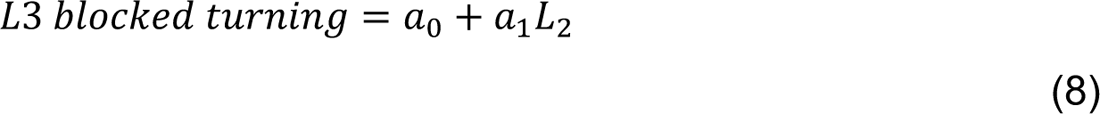 Where L_2_ denotes the contrast-sensitive LMC output as before (see Eq. (4)). The L3 contribution (control turning minus L3-blocked turning) was modeled as a sum of the dichotomous L3 contribution (Eq. (7)) and the edge inhibition. The latter was modeled as −b_3_ log *I_edge_* tanh *I_edge_*, where *I_edge_* denotes the luminance of the moving OFF edge and the *tanh* function tackles the limit when *I_edge_* approaches 0. Including this edge inhibition term, the total L3 contribution can be formulated as

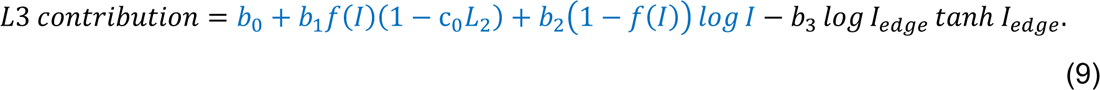 The predicted L3-blocked turning (Eq. (8)) and the predicted L3 contribution to turning (Eq. (9)) were then combined to predict turning under control conditions (Figure 7):

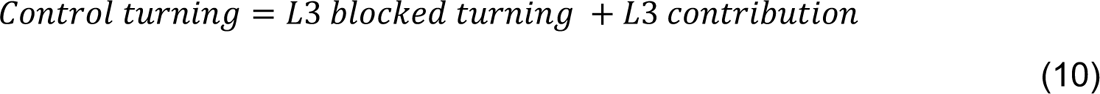 We first fitted the turning without gain control (Eq. (10)) on the L3-blocked fly turning of the uniform-contrast datasets, to obtain the two coefficients, *a_0_* and *a_1_*, together with the scaling constant of the L2 gain (*k_0_* in Eq. (5)). The remaining coefficients b_0_, b_1_, b_2_, b_3_ and c_0_ were obtained by fitting the L3 contribution using Eq. (9). For both equations, the fitting was performed in 30 iterations, with 50% of all the data points randomly chosen for training in each iteration, and the rest used for testing. Coefficient estimates obtained by fitting are shown in the Table S2. On the training set, the goodness of fit averaged over all iterations was *R^2^* = 0.904 for L3-blocked fly turning and *R^2^* = 0.908 for the L3 contribution (control turning – L3-blocked turning). On the testing set, the goodness of fit averaged over all iterations was *R^2^* = 0.861 combining the data from L3-blocked fly turning and from the control turning. If instead the predictions for the same stimulus from multiple iterations were first averaged and the goodness of fit of this average model was tested, the model exhibited a goodness of fit of *R^2^* = 0.873 on uniform-contrast datasets (Figure 7F). For separate contrasts, the goodness of fit of this averaged prediction was: *R^2^* = 0.850 (−50% uniform-contrast set), *R^2^* = 0.844 (−80% uniform-contrast set), and *R^2^* = 0.863 (−100% uniform-contrast set), respectively. We then used the fitted model to predict turning response to the mixed-contrast stimuli, generating 30 predictions in the 30 iterations. The average goodness of fit of these 30 predictions was *R^2^* = 0.854 (Figure 7G).
6. Model trained on mixed-contrast dataset To validate the model on an independent dataset, the same model (Eq. (10) and Eq. (9)) was separately trained on a mixed-contrast dataset and validated on the −50% and −80% uniform-contrast datasets (Figure S6). The −100% uniform-contrast dataset was excluded from this analysis, since the mixed-contrast data lacking −100% contrast would be insufficient to constrain the model to predict the predominant underestimation of −100% contrast by L3-blocked flies. As before, *a_0_*, *a_1_*, and *k_0_* were fitted on turning responses of L3-blocked flies using Eq. (10) and other coefficients were fitted on L3 contribution using Eq. (10), in 30 iterations. Coefficient estimates obtained by fitting are shown in the Table S3. On the training set, the goodness of fit averaged over all iterations was *R^2^* = 0.939 for L3-blocked responses and *R^2^* = 0.959 for the L3 contribution to control turning responses. On the testing set, the goodness of fit averaged over all iterations was *R^2^* = 0.822 combining the data from L3-blocked fly turning and from the control turning. If instead the predictions for the same stimulus from multiple iterations were first averaged and the goodness of fit of this average model was tested, the model exhibited a goodness of fit of *R^2^* = 0.854 on the mixed-contrast dataset. When predicting response to uniform contrasts datasets, the model performed with overall goodness of fit *R^2^* = 0.660 (−50% contrast: *R^2^* = 0.809, −80% contrast: *R^2^* = 0.659). For the further analyses involving the model, we obtained an ‘average model’ by averaging the scaling coefficients obtained over 30 iterations of fitting the model on the uniform-contrast datasets.
7. Sensitivity analysis of the model We performed a sensitivity analysis of the mixing coefficients in the model for the L3 contribution to total turning, i.e., *b*_j_ (j = 1,2,3) in Eq. (9). We changed each of the average coefficient values by a small percentage to quantify the model’s performance in predicting three aspects of the turning data: (A) underestimation and (B) overestimation of stimuli in L3-blocked fly turning, and (C) the dichotomy of the L3 contribution to control turning. To perform this sensitivity analysis, we first quantified the three aspects of the data as follows. For −100% contrasts, the underestimation was quantified as the average difference between the turning response of control and L3-blocked flies over all NDs (Figure S7A, red arrows). For −50%, −80%, and mixed contrasts, the underestimation was quantified as the same difference, but averaged only over the two dimmest adaptation contexts (ND 2.7 and ND 3.6). Overestimation was not defined for −100% contrasts because L3-blocked flies mostly only underestimate this contrast value. For −50%, −80%, and mixed contrasts, overestimation was quantified as the average difference between the turning response of control and L3-blocked flies over the two brightest adaptation contexts (ND 0 and ND 0.9, Figure S7B for −50% contrast, blue arrows). To quantify the dichotomy in the −100% contrast dataset, we first fitted the L3 contribution (from Figure 6A) with linear functions in separate segments (Figure S7C). We fitted four linear functions, three at luminances higher than the dichotomy point *I_m_* and one at luminances lower than *I_m_*. Denoting their slopes as *k*_1_, *k*_2_, *k*_3_, and *k*_4_, the dichotomy was then calculated as −(*k*_1_ + *k*_2_ + *k*_3_)*k*_4_/3. For −50%, −80%, and mixed contrasts, the dichotomy was defined as the product of underestimation and overestimation. If we denote one of these three quantities (underestimation, overestimation, and dichotomy) as *y*, the fitted coefficients in the average model as *^b_j_* (with *j* = 1,2,3), the relative change of coefficient *b_j_* as *z_j_* = (*b_j_* − *^b_j_*)/b|*^b_j_*~, then the sensitivity of *y* with respect to *b_j_* can be defined as

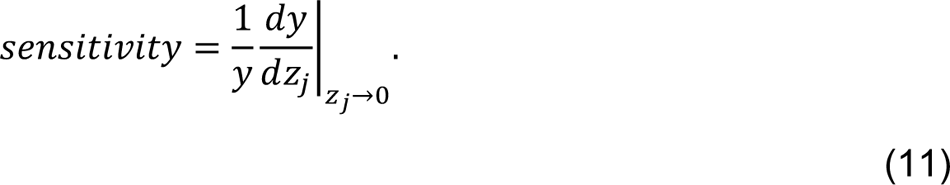 We calculated sensitivities at *z_j_* → 0 for all coefficients (Table S4). We illustrate how the model’s prediction with regards to underestimation, overestimation, and dichotomy changes when some of the average fitted coefficients were changed by *z_j_* = 0.1 (Figure S7,A’-C’ compared with A’’-C’’). For example, a change of 10% in b_1_ (the coefficient of f(*I*)(1 − c_&_L_2_) leads to a change of 34.2% in the predicted underestimation of contrasts for L3-blocked fly responses. A change of 10% in the coefficients b_1_c_0_ (the coefficient of −f(*I*)L_2_ and b_3_ (the coefficient of log *I_edge_* tanh *I_edge_*) respectively leads to a change of 12.4% and 12.9% in the predicted overestimation of contrasts for L3-blocked fly responses. Since the dichotomy in L3 contribution is primarily driven by the correction of underestimation at low luminances, the prediction of dichotomy also changes when a change of 10% in b_2_ (the coefficient of R1 − f(*I*)S log *I*) leads to a change of 23.7% in the predicted underestimation of dim stimuli by L3-blocked fly responses. The analysis further suggests that the model component with the coefficient b_1_implements both gain increments and decrements (Figure S7, A’,A’’ and B’,B’’) through the non-linear interaction of luminance and contrast channels. Differently, the coefficients b_2_ and b_3_ of the luminance-dependent model components separately implement either gain increments or decrements (Figure S7, B’,B’’ and C’,C’’). Overall, the distinct corrective terms in the model were involved in implementing specific types of gain control and all contributed to the model’s performance.
8. Predicted turning response assuming complete adaptation and the decomposition of L3 corrections Inserting *I_a_* = *I* into Eq. (2) results in *C_a_* = *C*. Then, based on Eq. (4), we could calculate the L2 response when it is fully adapted to the background luminance (Figure 8A):

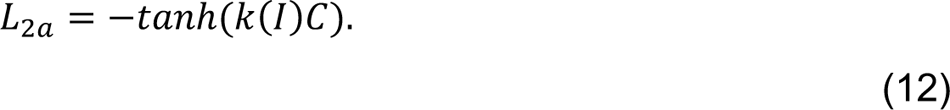 According to the model (Eq. (10)), the predicted turning response of L3-blocked flies under complete adaptation then is (Figure 8B, orange):

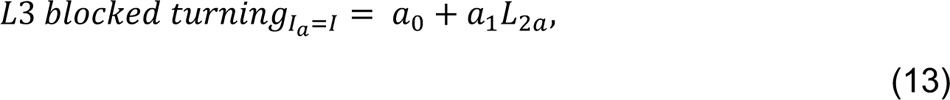

and the predicted turning response of control flies, according to Eq. (10), is (Figure 8B, blue):

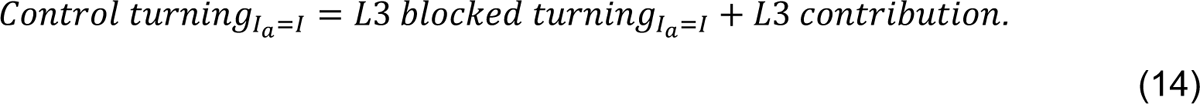 The predicted L3-blocked turning by Eq. (15) misses both fast- and slow-timescale gain correction required to achieve the control turning. However, the L3-blocked turning predicted by Eq. (16) misses only slow-timescale gain correction. Therefore, the fast-timescale correction can be obtained as the difference between the predicted L3-blocked fly turning by Eq. (17) and Eq. (18), which simplifies into *a_1_*[−tanh(*k*(*I*)*C*) − *L_2_*] (Figure 8D, bottom), while the slow-timescale correction is the difference between the predicted control turning by Eq. (19) and the predicted L3-blocked fly turning by Eq. (20), which simplifies into b_0_ + b_1_f(*I*)(1 − c_0_L_2_ + b_2_R1 − f(*I*)S log *I* − b_3_ log *I_edge_* tanh*I_edge_* (Figure 8B, bottom). Note that since *L_2a_* = −tanh(*k*(*I*)*C*) (Eq. (12)) and *I_edge_* = (1 + *C*)*I* (Eq. (3)), the slow-timescale correction can be written as b_0_ + b_1_f(*I*)[1 + c_0_ tanh(*k*(*I*)*C*)] + b_2_R1 −f(*I*)S log *I* − b_3_ log[(1 + *C*)*I*] tanh[(1 + *C*)*I*]. The average model, as stated above, was used to calculate the predictions in both experimental condition and under complete adaptation, as well as the fast-timescale and the slow-timescale corrections (Figure 8B-D).

## Acknowledgments

We thank Carlotta Martelli and Christopher Schnaitmann for comments on the manuscript and are grateful to Jan Benda for informative discussions on adaptation. This project has received funding from the German Research Foundation (DFG) through the Emmy-Noether program (SI 1991/1-1), the collaborative research center 1080 “Neural homeostasis” (project C06) to MS and (project C07) to JG and the Max Planck Society (SS and JG).

## Supplemental Information

### 1. Supplemental figures

**Figure S1:**
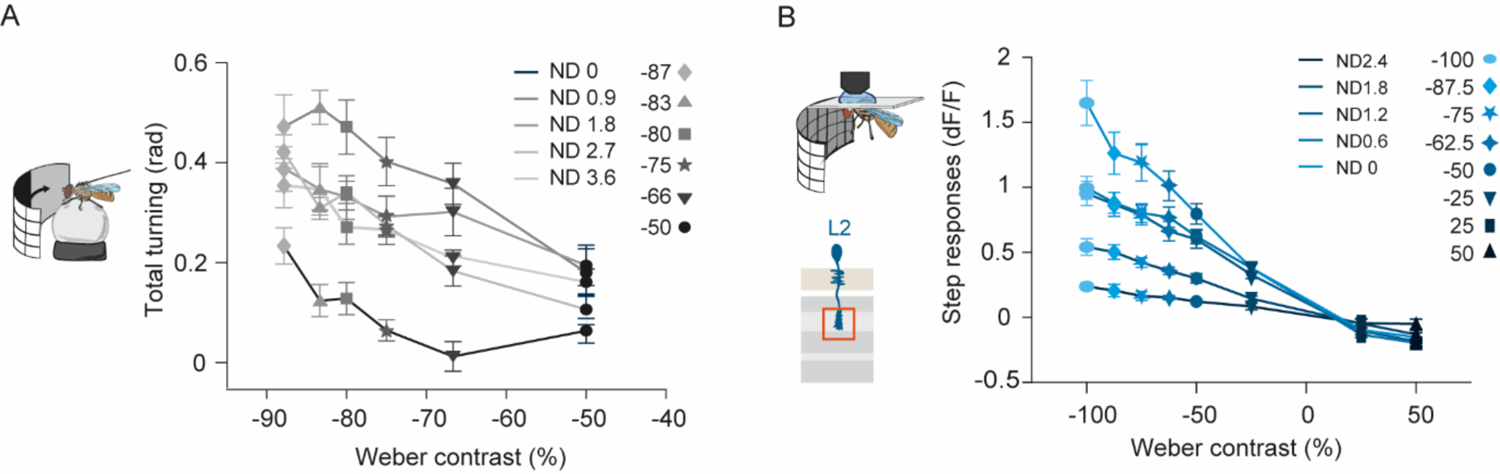
Contrast sensitivity of turning behavior and L2 calcium responses are fundamentally different. (A) Quantification of total turning in response to different contrasts, in five adaptation contexts (denoted by the ND filter) (same data as in Figure 2B). Sample size in n=10 per data point. (B) L2 calcium response to contrast steps quantified as the difference between peak response and preceding plateau response in five adaptation contexts (same data as in Figure 2D). Sample size is n=10 (136), #flies (#ROIs). Both plots show mean ± SEM.

**Figure S2:**
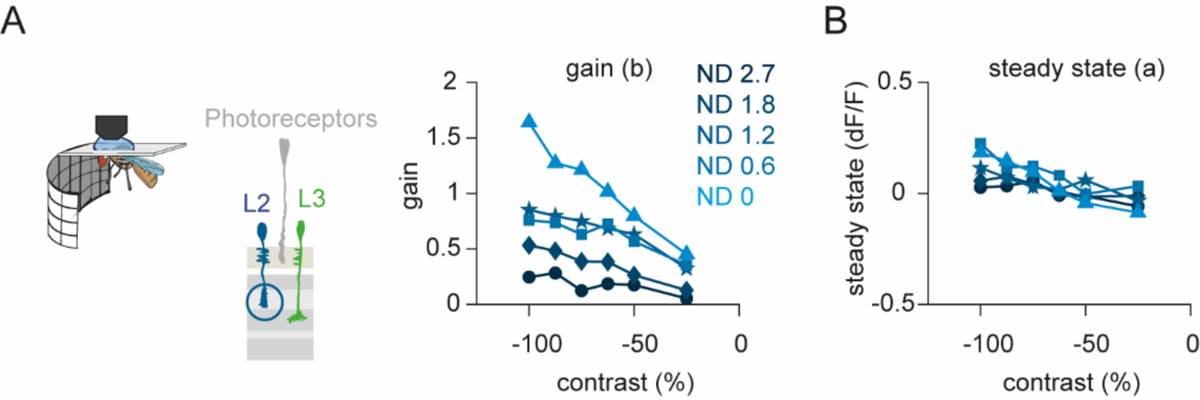
Properties of L2 adaptation. (A) Exponential functions (L2 response = *a + b * exp(−c * time)*) were fit to the L2 calcium responses (data from Figure 2D). The gain (b) of the exponential fits is plotted for all negative contrasts tested in five adaptation contexts. (B) The steady-state parameter (*a*) of the exponential fits. In both plots, the blue shades represent adaptation context (also denoted by the ND filter). Data are shown as mean ± SEM. Sample size is n=10 (136), #flies (ROIs).

**Figure S3:**
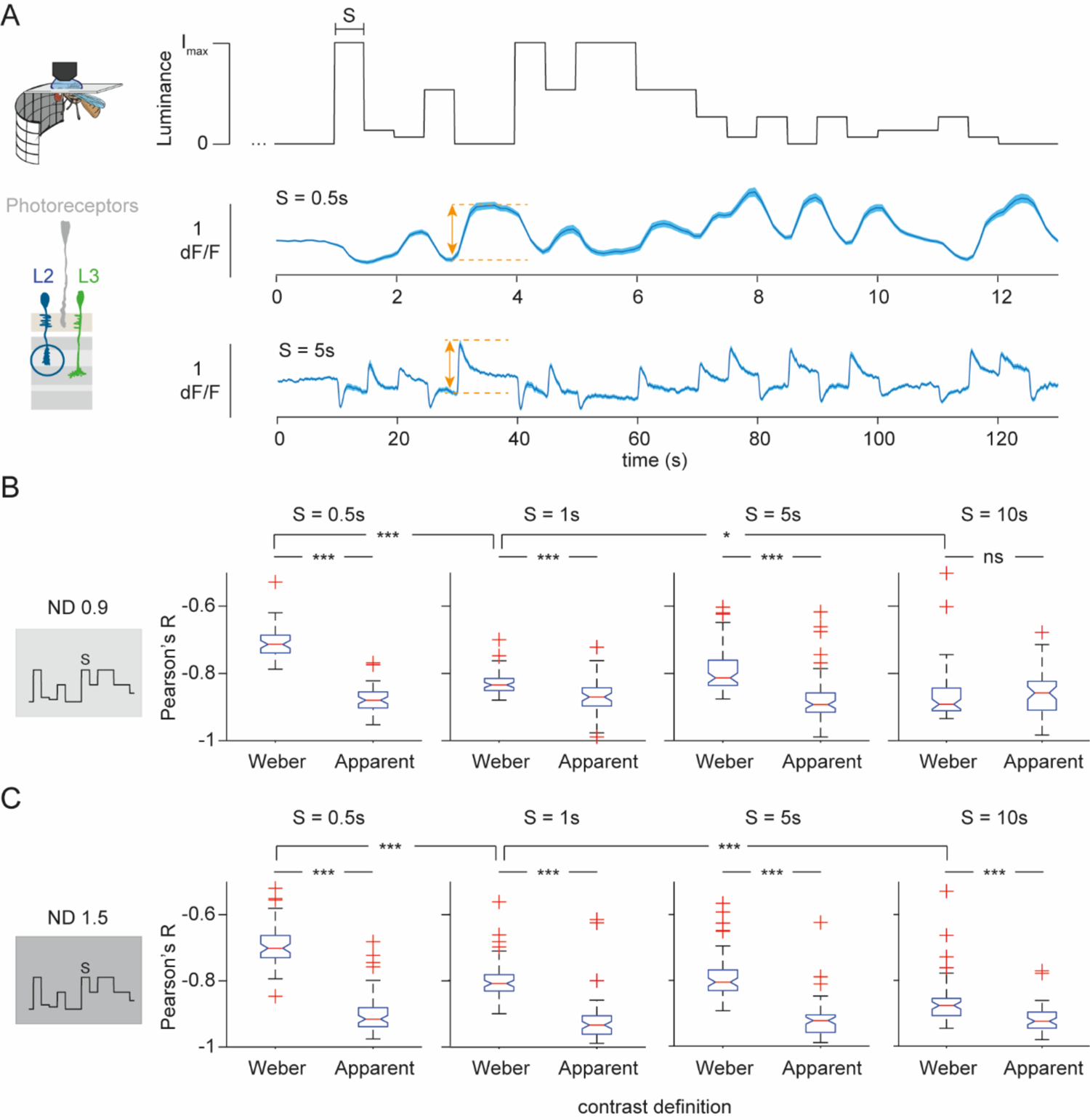
L2 responses correlate stronger with apparent contrast than with Weber contrast. (A) L2 calcium signals in response to random luminance steps. (Top) An example stimulus trace. The stimulus comprised a sequence of full-field flashes, where luminance stepped randomly between six values (0%, 7%, 14%, 27%, 53% and 100% of I_max_. ND 0.9: I_max_=3.06*10^4^ photos s^-1^ receptor^-1^, ND 1.5: I_max_=7.68*10^3^ photos s^-1^ receptor^-1^). The step durations (S) 0.5s, 1s, 5s and 10s were tested in separate experiments. L2 calcium signals in response to 0.5s luminance steps (middle, n=52 ROIs) and 5s luminance steps (bottom, n=65 ROIs) are plotted as mean ± SEM. The dashed orange lines mark the pre-step and post-step calcium signals, and their difference is quantified as L2 step response (orange arrows). (B-C) Coefficients of correlation (Pearson’s R) between the L2 step response and the luminance step stimuli, quantified as either Weber contrast or apparent contrast (Figure 4A), and the L2 step responses elicited by them. Note that L2 responses negatively correlate with luminance changes, and thus a more negative R indicates a higher correlation. (B) R values in a relatively bright adaptation context (ND 0.9). For the three smaller values of S (0.5s, 1s and 5s), L2 responses correlate more strongly with apparent contrasts than with Weber contrasts. n=52, 52, 65 and 51 ROIs for S=0.5s, 1s, 5s and 10s respectively. (C) R values in a relatively dim adaptation context (ND 1.5). For all values of S, L2 responses correlate more strongly with apparent contrasts than with Weber contrasts. n=64, 64, 63 and 55 ROIs for S=0.5s, 1s, 5s and 10s respectively. (B,C) When only comparing the correlation with Weber contrast at different times S, the correlation is stronger at S=1s than at S=0.5s, and stronger at S=10s than at S=1s, in line with the long adaptation time constants of L2. Boxplots in (B) and (C) show median and 25^th^ and 75^th^ percentile, notches mark 5% comparison intervals and whiskers extend to the most extreme data points excluding outliers (red ‘+’). Paired sample t-tests corrected for multiple comparisons were used to compare response correlation with Weber contrast and apparent contrast; two-sample Student t-tests corrected for multiple comparisons were used to compare response correlation with Weber contrasts between different S values. *p<0.05, ***p<0.001, ns = non-significant.

**Figure S4:**
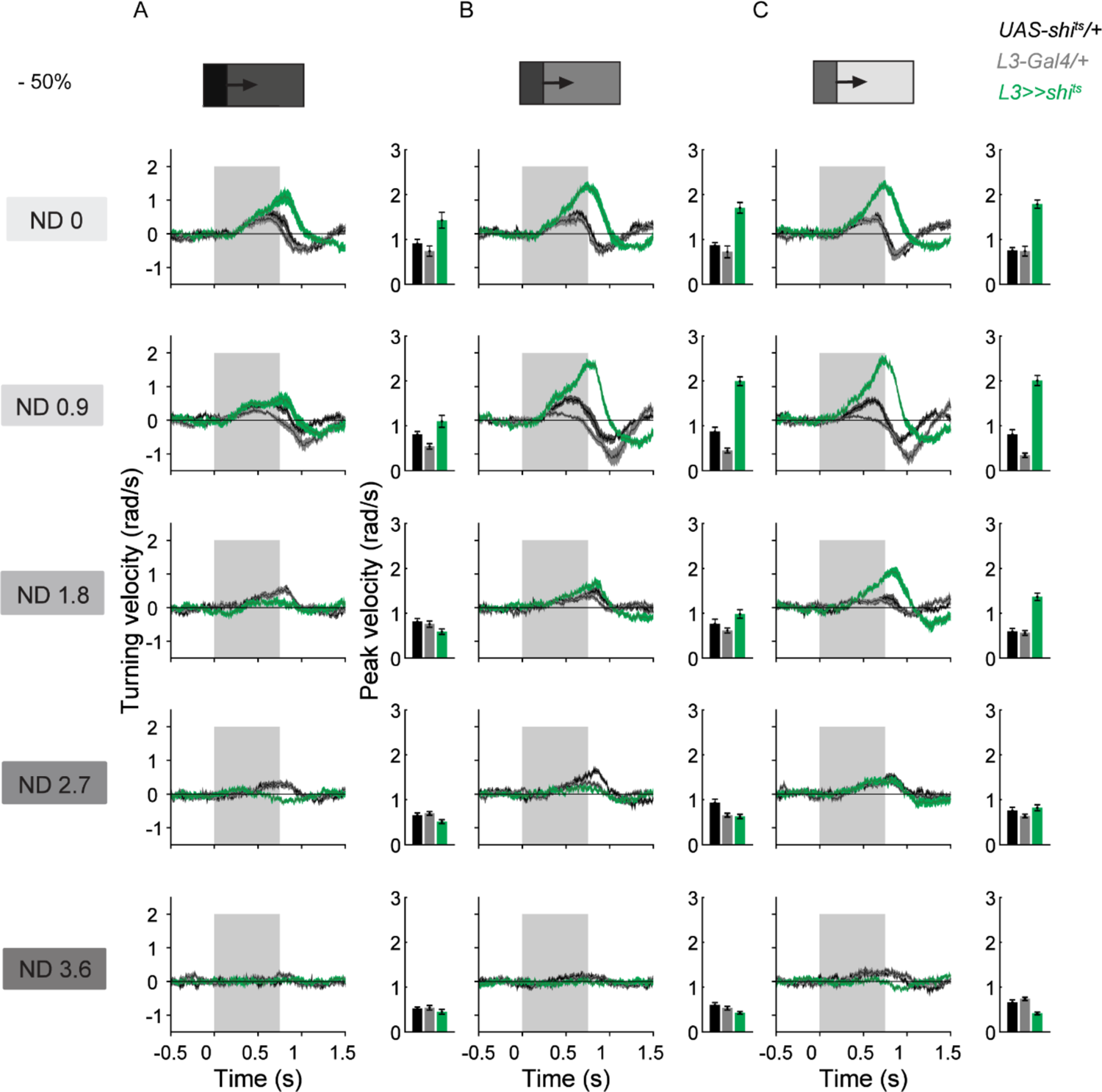
At −50% contrast, response of L3-blocked flies underestimates stimuli in low adaptation contexts and overestimates them in the bright adaptation context. (A-C) Time traces and peak turning velocities of L3-blocked flies (*L3>>shi^ts^*, green) and their genetic controls (*L3-Gal4/+ and UAS>>shi^ts^*, gray and black) to the uniform-contrast stimuli at −50% contrast, same dataset as in Figure 4E. The columns represent three of the five edges moving onto different background luminances, within each adaptation context: the edge with (A) the dimmest, (B) an intermediate and (C) the brightest contextual luminance. Rows correspond to five different adaptation contexts, generated using ND filters. Sample size in n=10 per plot. Traces and plots show mean ± SEM.

**Figure S5:**
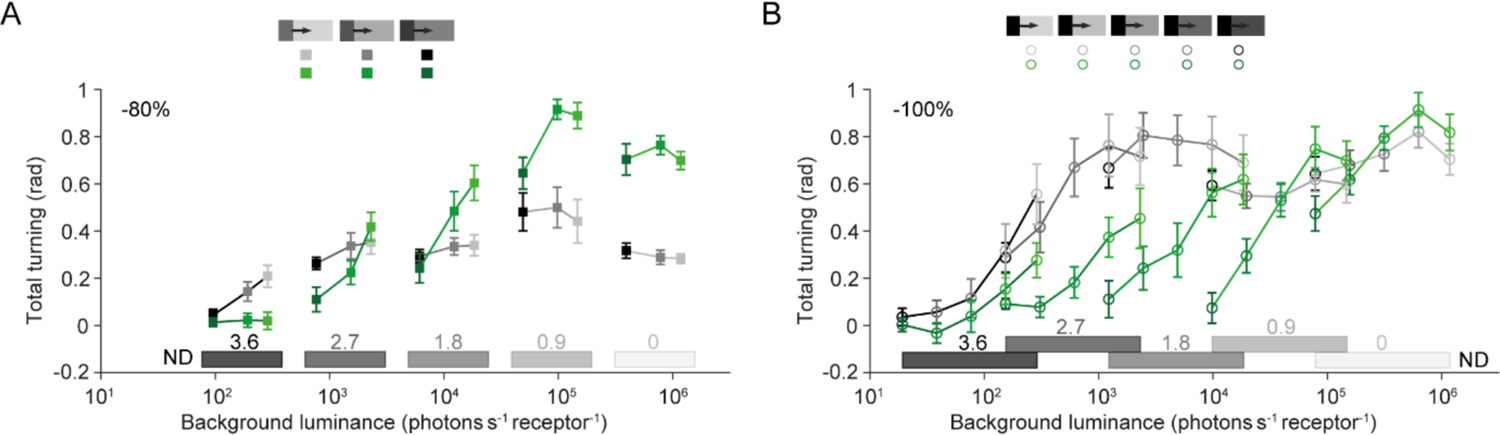
Turning response of control and L3-blocked flies to uniform-contrast stimuli in five adaptation contexts. (A) Total turning in response to −80% contrast moving edges, by control files (gray) and L3-blocked files (green). (B) Same as in A, but for −100% contrast moving edges, re-quantified as total turning from the data published in (4). Sample size is n=10 per plot. Both plots show mean ± SEM.

**Figure S6:**
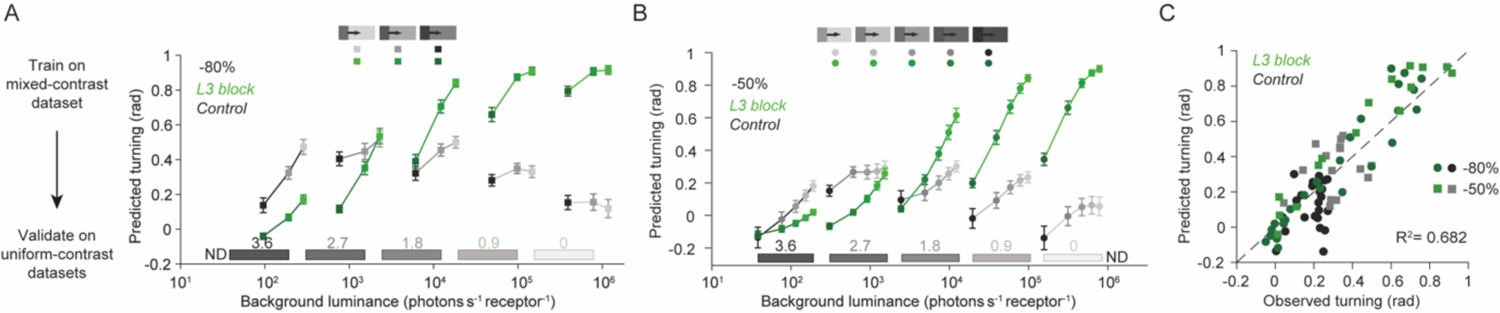
Additional cross-validation of the model. (A) Predicted turning behavior of control and L3-blocked flies in response to −80% contrast stimuli as predicted by the model fitted on mixed-contrasts data. (B) Same as A, but for −50% contrast. Gray: control flies; green: L3-blocked flies. Both plots show mean ± SD over 30 iterations. (C) Mean predicted turning responses from A and B plotted against the corresponding observed turning responses (Figure 4E, S5A).

**Figure S7:**
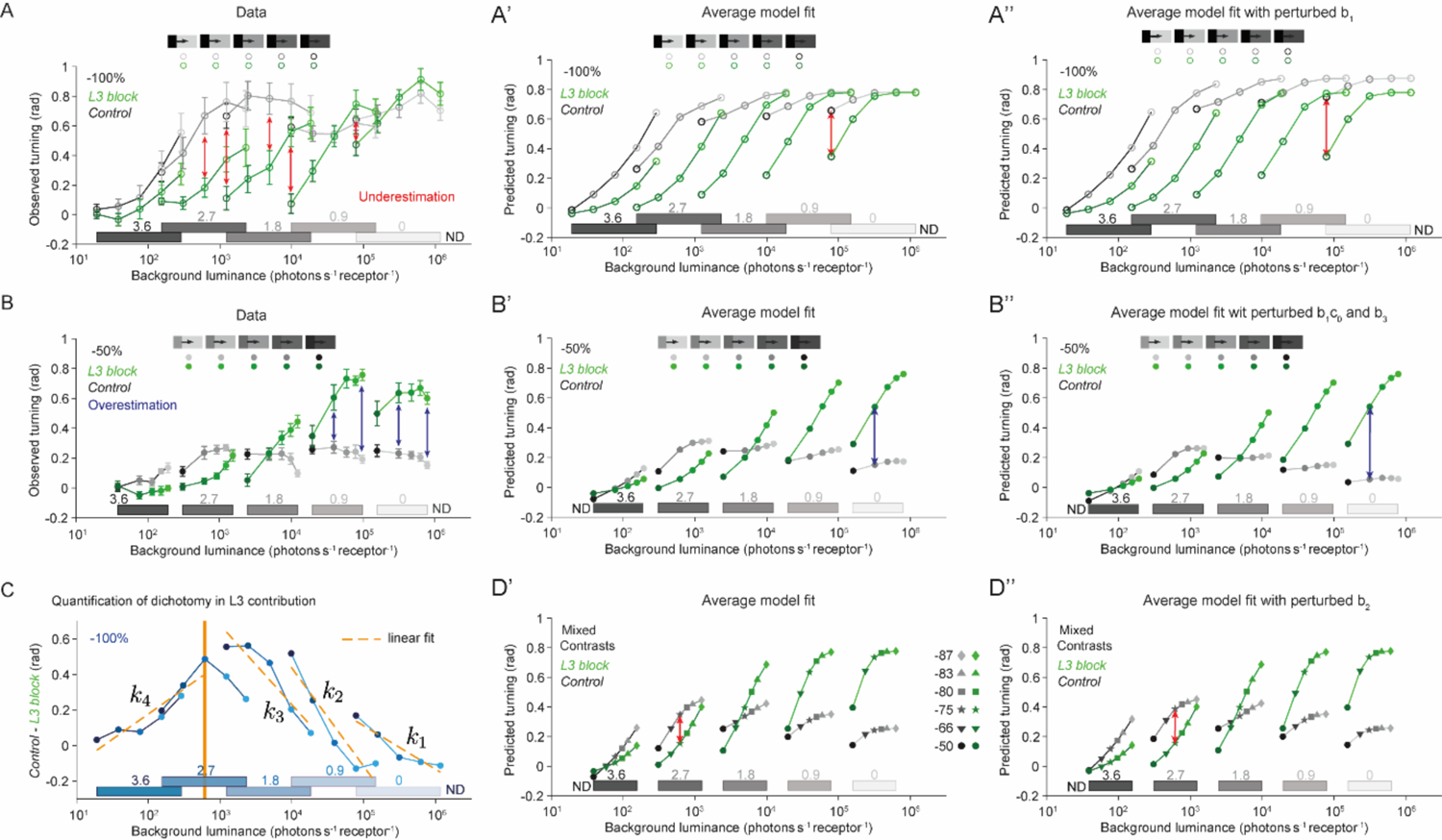
Sensitivity analysis of the model parameters reveals specific gain control functions of different components of the model. The model’s performance tested in the sensitivity analysis was quantified for three different data features: (A) Underestimation and (B) overestimation of stimuli from L3-blocked fly turning responses, and (C) dichotomy in L3 contribution to total turning. Prediction of these features was compared under two modeling conditions: (A’-D’) predicted total turning using coefficients of the average model (see Methods) and (A’’-D’’) predicted turning when specific coefficients of the average model were changed by 10%. (A) Turning response of control and L3-blocked flies to −100% contrast. The red arrows point at the underestimation of some of the stimuli by L3-blocked flies. Data are from (4), quantified as total turning (also see Figure S5B). (A’,A’’) The model differently predicts underestimation of −100% in bright conditions when the coefficient b_1_ of the model component f(*I*)(1 − c_0_L_2_) is increased by 10%. (B) Turning response of control and L3-blocked flies to −50% contrast. The blue arrows point at the underestimation of some of the stimuli by L3-blocked flies (also see Figure 4E). (B’,B’’) The model differently predicts overestimation of −50% in bright conditions when the coefficients b_1_c_$_ (corresponding to the model component-*f(1)(L_2_)*) and *b_3_* (corresponding to the model component log *I*_edge_ tanh *I*_edge_) are both increased by 10%. (C) The difference between the turning response of control and L3-blocked flies to −100% contrast (same as Figure 6A), overlaid with piecewise linear fits over different luminance ranges (see Methods). The product of the slopes of the linear fits on the two sides of the dichotomy point was used as the measure of the dichotomy (see Methods). (D’,D’’) The model differently predicts underestimation of −75% contrast (from the mixed-contrast experiment) in dim conditions when *b_%_* (coefficient of the model component B1 − f(*I*)C log *I*) is increased by 10%. In (A-B) and (A’-D’, A’’-D’’), control flies: gray, L3-blocked flies: green.

**Figure S8.**
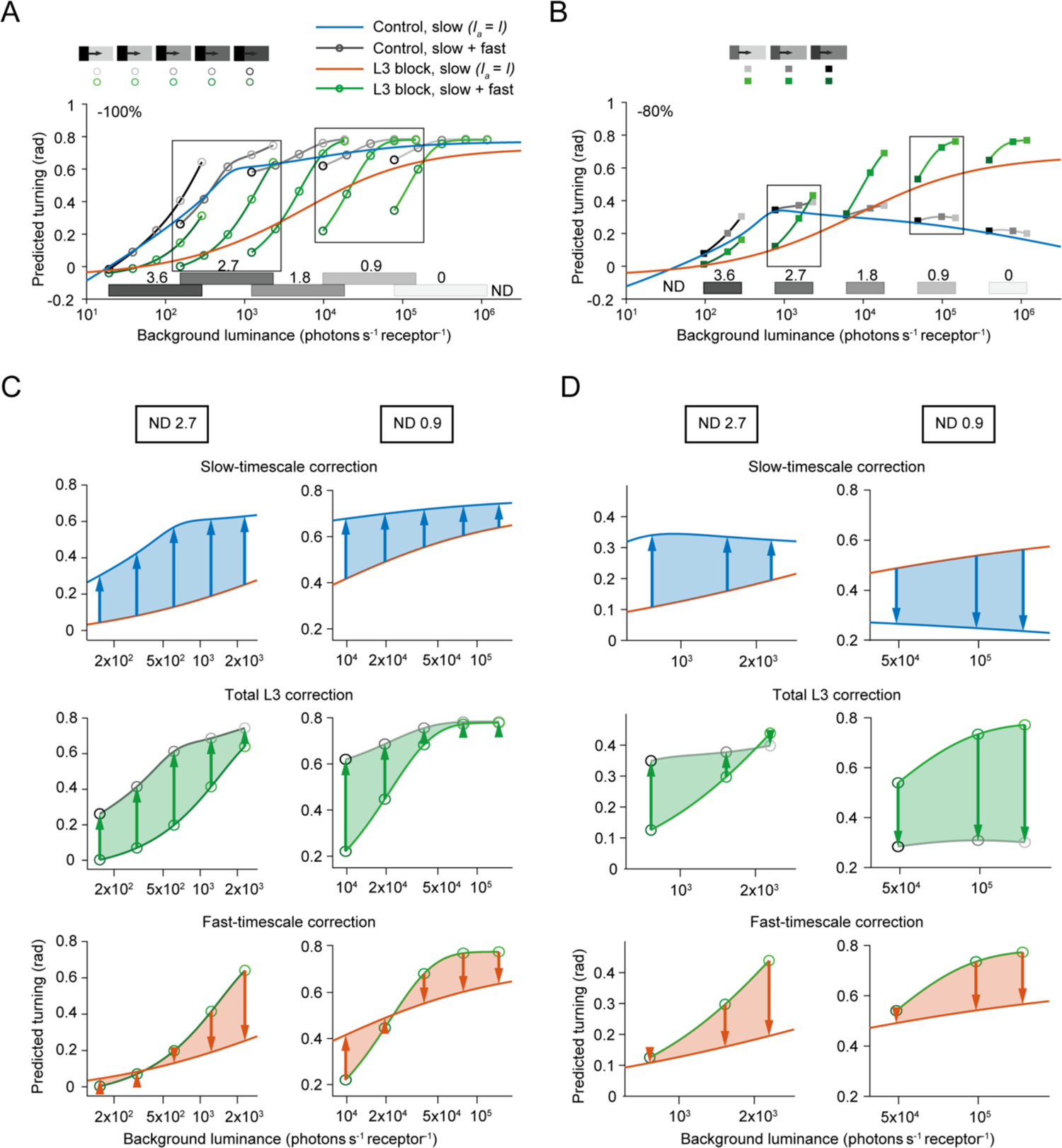
The decomposition of the corrective mechanism of L3 for −100% and −80% uniform contrast stimuli. (A) Predicted turning response to −100% contrast by the average model, for control files (gray) and L3-blocked files (green), at slow + fast timescales, superimposed onto the predictions at slow timescales where L2 fully adapts (blue for control and orange for L3-blocked flies). Boxes indicate zoomed-in intervals in C. (B) Same as A, but for −80% OFF contrast. Boxes indicate the zoomed-in intervals in D. (C,D) The corrective contribution of L3 at −100% (C) and −80% (D) contrasts decomposed into the fast-timescale correction and the slow-timescale correction. Left: ND 2.7 (the box on the left in A). Right: ND 0.9 (the box on the right in A). Top: The slow-timescale L3 correction illustrated as the discrepancy in the predicted turning response between control and L3-blocked flies when *I* = *I_a_*. Middle: Total L3 correction illustrated as the discrepancy between the predicted control and L3-blocked fly response, both under experimental condition (slow + fast timescales). Bottom: The fast-timescale L3 correction illustrated as the discrepancy between the predicted L3-blocked fly responses under experimental condition (fast + slow timescales) and under the hypothesized complete adaptation condition (slow timescales, *I* = *I_a_*). The arrows point in the direction of gain control: upwards for gain increment, downwards for gain decrements.

### 2. Supplemental tables

**Table S1:** Linear Model fitted on −100% contrast data (Figure 4D) – parameter estimates Coefficient for L2 gain *k_0_* = 1.26.

| Coefficient | Estimate | SE | p-Value |
| --- | --- | --- | --- |
| $a_0$ | -8.1E-4 | 0.028 | 0.98 |
| $a_1$ | 0.781 | 0.047 | 2.4E-14 |

Goodness of fit: $R^2 = 0.924, p = 2.4 \times 10^{-14}$ .

**Table S2:** Model (Eq. (10)) fitted on the uniform-contrast data (Figure 7) – parameter estimates and goodness of fit.

| Coefficient | Mean | SD | p-Value (t-test) |
| --- | --- | --- | --- |
| $k_0$ | 1.785 | 0.139 | 5.3E-34 |
| $a_0$ | -0.064 | 0.015 | 2.0E-20 |
| $a_1$ | 0.842 | 0.029 | 5.2E-44 |
| $b_0$ | -0.312 | 0.077 | 1.0E-19 |
| $b_1$ | 0.912 | 0.075 | 3.3E-33 |
| $b_1 c_0$ | 0.594 | 0.040 | 1.0E-35 |
| $b_2$ | 0.261 | 0.037 | 1.7E-26 |
| $b_3$ | 0.108 | 0.008 | 4.0E-34 |

Goodness of fit on the training set, averaged over all iterations: $R^2 = 0.904$ for L3 blocked responses and $R^2 = 0.908$ for L3 contribution.
On the testing set, averaged over all iterations: $R^2 = 0.861$ .
Average predictions on uniform contrasts: $R^2 = 0.873$ .
Average predictions on -100% contrast: $R^2 = 0.863$ .
Average predictions on -80% contrast: $R^2 = 0.844$ .
Average predictions on -50% contrast: $R^2 = 0.850$ . Average predictions on mixed contrasts: $R^2 = 0.876$ .

**Table S3:** Model (Eq. (10)) fit on the mixed-contrast data – parameter estimates

| Coefficient | Estimate | SD | p-Value (t-test) |
| --- | --- | --- | --- |
| $k_0$ | 2.002 | 0.245 | 2.53E-28 |
| $a_0$ | -0.152 | 0.025 | 1.27E-24 |
| $a_1$ | 1.068 | 0.033 | 2.13E-45 |
| $b_0$ | -0.495 | 0.159 | 1.28E-16 |
| $b_1$ | 1.352 | 0.173 | 9.69E-28 |
| $b_1c_0$ | 0.375 | 0.107 | 4.90E-18 |
| $b_2$ | 0.494 | 0.077 | 2.13E-25 |
| $b_3$ | 0.238 | 0.034 | 2.85E-26 |

1331 Goodness of fit on the training set, averaged over all iterations: $R^2 = 0.939$ for L31332 blocked responses and $R^2 = 0.959$ for L3 contribution.1333 On the testing set, averaged over all iterations: $R^2 = 0.822$ .1334 Average predictions on mixed contrasts: $R^2 = 0.854$ .1335 Average predictions on -50% and -80% contrasts: $R^2 = 0.682$ .1336 Average predictions on -80% contrast: $R^2 = 0.659$ .1337 Average predictions on -50% contrast: $R^2 = 0.660$ .

**Table S4:** The results of sensitivity analysis

|  | Underestimation |  |  |  |  |
| --- | --- | --- | --- | --- | --- |
|  | -100% | -80% | -50% | Mixed | Average |
| $a_0$ | 0.000 | 0.000 | 0.000 | 0.000 | 0.000 |
| $a_1$ | 0.000 | 0.000 | 0.000 | 0.000 | 0.000 |
| $b_0$ | 1.672 | 3.198 | 3.700 | 3.363 | 2.983 |
| $b_1$ | 3.415 | 4.272 | 3.613 | 2.704 | 3.501 |
| $b_1c_0$ | -1.612 | -1.204 | -0.609 | -0.694 | -1.030 |
| $b_2$ | 0.869 | 3.331 | 4.445 | 4.368 | 3.253 |
| $b_3$ | 0.000 | -2.202 | -2.748 | -2.014 | -1.741 |

|  | Overestimation |  |  |  |  |
| --- | --- | --- | --- | --- | --- |
|  | -100% | -80% | -50% | Mixed | Average |
| $a_0$ | -- | 0.000 | 0.000 | 0.000 | 0.000 |
| $a_1$ | -- | 0.000 | 0.000 | 0.000 | 0.000 |
| $b_0$ | -- | -0.691 | -0.879 | -0.826 | -0.799 |
| $b_1$ | -- | -2.019 | -2.566 | -2.413 | -2.333 |
| $b_1c_0$ | -- | 1.204 | 1.212 | 1.318 | 1.244 |
| $b_2$ | -- | 0.000 | 0.000 | 0.000 | 0.000 |
| $b_3$ | -- | 1.123 | 1.476 | 1.268 | 1.289 |

|  | Dichotomy |  |  |  |  |
| --- | --- | --- | --- | --- | --- |
|  | -100% | -80% | -50% | Mixed | Average |
| $a_0$ | 0.000 | 0.000 | 0.000 | 0.000 | 0.000 |
| $a_1$ | 0.000 | 0.000 | 0.000 | 0.000 | 0.000 |
| $b_0$ | 0.000 | 2.506 | 2.821 | 2.537 | 1.966 |
| $b_1$ | 0.709 | 2.254 | 1.047 | 0.291 | 1.075 |
| $b_1c_0$ | 0.854 | 0.000 | 0.602 | 0.624 | 0.520 |
| $b_2$ | 0.436 | 3.331 | 4.445 | 4.368 | 3.145 |
| $b_3$ | 0.000 | -1.079 | -1.272 | -0.746 | -0.774 |

